# Beta-DIA: Integrating learning-based and function-based feature scores to optimize the proteome profiling of single-shot diaPASEF mass spectrometry data

**DOI:** 10.1101/2024.11.19.624419

**Authors:** Jian Song, Hebin Liu, Chengpin Shen, Xiaohui Wu

## Abstract

We present a freely available diaPASEF data analysis software, Beta-DIA, that utilizes deep learning methods to score coelution consistency in retention time-ion mobility dimensions and spectrum similarity. Beta-DIA integrates these learning-based scores with traditional function-based scores, enhancing the qualitative analysis performance. In some low detection datasets, Beta-DIA identifies twice as many protein groups as DIA-NN. The success of Beta-DIA has paved another way for the application of deep learning in fundamental proteome profiling.

## 1 Introduction

Liquid chromatography coupled with mass spectrometry (LC-MS) has become increasingly successful in the in-depth characterization of complex proteomes. Common mass spectrometry acquisition methods include data dependent acquisition (DDA) and data independent acquisition (DIA). Compared to DDA, which randomly samples and fragments high-intensity peptide ions, the DIA method employs consecutive scanning over predefined isolation windows along the mass-to-charge (*m/z*) dimension, enabling unbiased sampling of peptide ions for fragmentation^[1]^. DIA records fragmentation signals from peptide ions more comprehensively than DDA (*i.e.*, a higher ion sampling rate), offering the potential to identify more peptides/proteins from mass spectrometry data^[2]^. The narrow-window DIA method, utilizing the Astral mass spectrometer, reduces the DIA scan window to 2Th without increasing the cycle time, significantly simplifying the DIA spectra and greatly enhancing data sensitivity^[3]^.

However, the DIA method has a limitation in ion sampling efficiency, failing to generate MS/MS spectra for all peptide ions present in a sample to the fullest extent. This is because peptide ions outside the isolation windows during a DIA scan cycle are essentially discarded. For instance, in typical 32- or 64-windows DIA methods, the corresponding ion sampling rates are only about 3.1% and 1.6%, respectively^[4]^. A method to improve the ion sampling rate of DIA without increasing spectral complexity is diaPASEF on timsTOF series mass spectrometers^[4]^. This technique achieves parallel accumulation-serial fragmentation (PASEF) by integrating two trapped ion mobility spectrometers (TIMS) between the chromatography and the mass spectrometer^[5]^. When using a typical four-scan diaPASEF scheme, it improves the ion sampling efficiency by fivefold compared to the standard DIA method. In addition to significantly enhancing ion sampling rate, diaPASEF expands the dimensionality of ion signal acquisition from three dimensions (retention time RT, *m/z* and intensity) to four (RT, ion mobility, *m/z* and intensity), further increasing the specificity of ion signals. These features have advanced the application of diaPASEF in fields such as single-cell proteomics^[6]^ and plasma proteomics^[7]^. However, these advantages also present challenges for diaPASEF analysis software. Effectively utilizing the four dimensions of diaPASEF data and converting its high sampling rate into more peptide/protein identifications are fundamental issues that diaPASEF analysis software needs to address.

The identification strategy of diaPASEF is derived from that of DIA data, and can be categorized into spectrum-centric and peptide-centric methods^[8]^. The latter is considered to offer greater detection sensitivity, as it more effectively incorporates RT, ion mobility and fragmentation patterns provided by spectral libraries. Existing software supporting diaPASEF, such as OpenSWATH^[4, 9]^, MaxDIA^[10]^ and DIA-NN^[11, 12]^, all adhere to the peptide-centric identification paradigm. The logic of this paradigm is as follows: For each target peptide from a spectral library that needs to be interrogated, it first assumes the peptide is existence. Then, it characterizes or quantifies the evidence or features of the existence across multiple aspects (such as RT, MS and MS/MS). Finally, it evaluates the relative strength of these features to confirm or reject the initial hypothesis, thereby completing the identification process. Clearly, feature quantification is the most critical step of peptide-centric DIA identification. Different DIA analysis software use various methods to quantify features. OpenSWATH developed 23 feature quantification functions, covering the co-elution consistency calculation of fragment ions in both the RT and ion mobility dimensions. MaxDIA introduced 60 subscores, including the deviation between ions’ actual and estimated mobility values. DIA-NN proposed a 2D-peak-picking algorithm that transforms the diaPASEF data from profile into centroid format, allowing the direct application of 73 scoring functions designed for DIA data. Several evaluation studies have shown that DIA-NN’s feature design results in superior qualitative outcomes, and combined with its outstanding quantitative capabilities, processing speed, and ease of use, it has established itself as the de facto standard for the free analysis of DIA and diaPASEF data, as well as a core infrastructure for the reanalysis of public large-scale proteomics data^[13, 14]^.

However, the feature scoring in the aforementioned analysis software relies on specific mathematical functions selected by manual experience. When it comes to quantifying complex features, these function-based scoring methods struggle to effectively mitigate the impact of noise, leading to inaccurate feature scores and suboptimal identification performance. For instance, in DIA data, the consistency of the fragment ions’ elution profiles, known as coelution consistency^[15]^, is not only the primary feature used to determine the presence of target peptide ions but also one of the most complex features to quantify. Elution profiles involve signals from multiple fragment ions at multiple time points, making them highly susceptible to signal loss, shifts, and distortions. Although various functions have been employed in the above-mentioned software to quantify coelution consistency, including dot product, cross-correlation^[15]^, cosine similarity^[16]^, peak shape similarity^[17]^, and entropy functions^[18]^, they still fail to overcome these interferences, resulting in inaccurate consistency calculation. Alternatively, deep learning or data-driven scoring offers another approach to feature quantification. Alpha-XIC^[19]^ and Alpha-Tri^[20]^ were the first to introduce learning-based scoring methods that compute the coelution consistency and spectrum similarity with greater accuracy and robustness, thereby aiding DIA-NN in identifying more peptides/proteins from DIA data. DreamDIA^[21]^ took this a step further by directly incorporating deep learning feature layers as scoring items, leading to additional gains in identification performance. These findings demonstrate that some complex features of DIA data are better suited for quantification through data-driven methods. As DIA data evolves into diaPASEF data, the increased dimensionality further complicates the data structure of features. Given the powerful information extraction capabilities of deep learning, it is reasonable to believe that learning-based scoring can more accurately quantify certain complex features of diaPASEF data compared to function-based scoring, thereby further enhancing detection sensitivity.

This study extends the concept of learning-based scoring for diaPASEF data by introducing a new analysis software, Beta-DIA. Its feature scoring combines learning-based and function-based methods. The learning-based scoring focuses on quantifying complex two-dimensional (RT and ion mobility) coelution consistency and spectrum similarity in diaPASEF data, while the function-based scoring leverages existing scoring functions to quantify other relatively simple features. Extensive testing data of diaPASEF from diverse samples, throughput, loading amounts and mass spectrometry instruments demonstrate that Beta-DIA can achieve a higher peptide/protein identifications compared to DIA-NN and provides accurate relative quantification results. In some low detection datasets, Beta-DIA identifies twice as many protein groups as DIA-NN. Beta-DIA, implemented by Python, is freely available and delivers high-sensitivity proteomics data support for downstream biological applications based on the diaPASEF method.

## 2 Results

### 2.1 The Overview of Beta-DIA

As shown in Figure 1, Beta-DIA performs a peptide-centric identification process using the input from a spectral library and .d files. Specifically, Beta-DIA extracts chromatograms for each target or decoy peptide ions from the centroided diaPASEF data and identifies potential elution groups. Beta-DIA then scores each elution group using both function-based and learning-based methods. Function-based scores include coelution consistency in RT dimension, the deviation between ions’ actual and estimated RT, mass accuracy, mobility accuracy and so on. Learning-based scoring is achieved through deep learning models DeepProfile (see Supplementary Figure 1) and DeepMall (see Supplementary Figure 2). DeepProfile scores coelution consistency in 2D (RT and mobility) dimensions, operating on the 2D elution groups which are retrieved from profile data (2D elution groups are produced from four-dimensional data cuboids by maximizing along the *m/z* dimension, preserving both chromatograms and mobilograms information). DeepMall computes the spectrum similarity between the measured and estimated intensities of fragment ions, weighting each ion based on various factors such as mass and mobility accuracy. Detailed descriptions of all scoring items can be found in Supplementary Table 2.

**Fig. 1.**
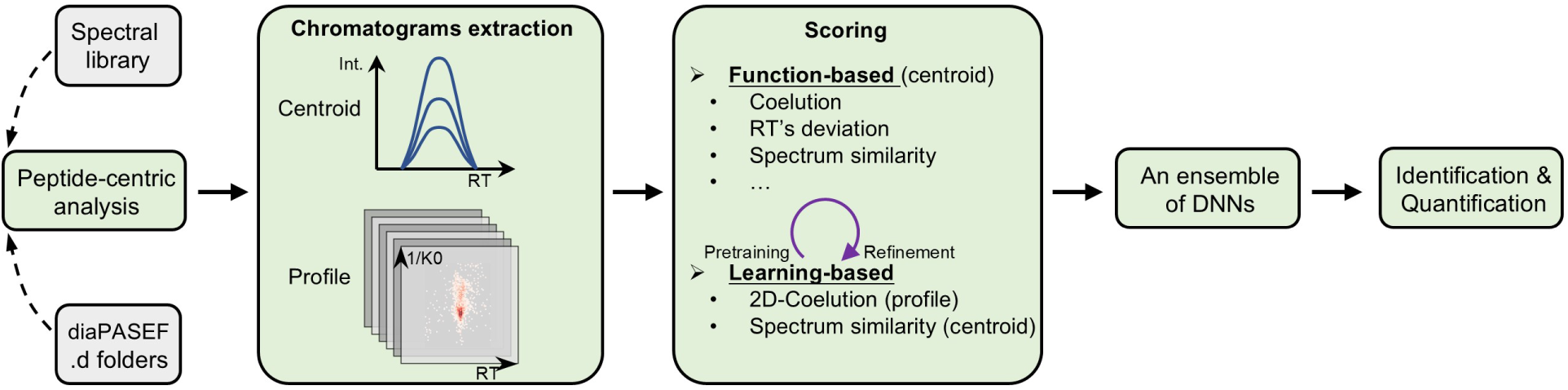
The schematic of Beta-DIA workflow. Beta-DIA extracts potential elution groups from centroid MS data and the corresponding 2D (RT-mobility) elution groups from profile MS data for each target or decoy peptide. It then applies both function-based and learning-based scoring, and aggregates these scores using an ensemble of deep neural networks (DNN) to calculate q-values. Finally, Beta-DIA generates a report with both qualitative and quantitative results.

After completing both function-based and learning-based scoring, Beta-DIA uses an ensemble neural network to distinguish between target and decoy peptides based on these scores and calculates the false discovery rate (FDR). Following peptide identification, Beta-DIA further realizes the protein inference and quantification.

### 2.2 Datasets

Since the implementation of Beta-DIA relies on a pretrained DeepProfile model, and it is impossible to encompass all potential interferences of 2D elution groups during pretraining (see Supplementary Table 3 for detailed pretraining data), a comprehensive assessment of Beta-DIA’s feasibility and qualitative/quantitative performance requires extensive testing with diverse data. This includes employing different diaPASEF cycle schemas, varying sample types and load amounts, using different mass spectrometers, and testing across various throughput levels. Supplementary Table 1 outlines the test data used in this study. Among these, the Base-X data originates from datasets released during the development of the diaPASEF method^[4]^ and is also used by DIA-NN to evaluate its performance^[12]^. Base-1 and Base-2 are utilized to verify Beta-DIA’s adaptability to different diaPASEF cycle schemas. Base-3 represents high-throughput data, Base-4 is a dilution series data, and Base-5 is two-proteome quantitative benchmark data. Besides, we add three additional datasets. Extra-1 provides higher throughput data, which includes ultra-high-throughput data (Extra-1-1, 400 SPD, SPD = samples per data) and high-throughput two-proteome quantitative data (Extra-1-2)^[22]^. Extra-2 is plasma sample data^[7]^ on timsTOF HT, and Extra-3 is single-cell data^[6]^ on timsTOF SCP (Extra-3-1) and Ultra (Extra-3-2).

### 2.3 Performance

We compared the performance of Beta-DIA and DIA-NN across three key aspects: qualitative performance, quantitative performance, and analysis efficiency. The qualitative performance metrics include the number of identifications per run and data completeness across triplicate samples. The quantitative performance metrics encompass the precision and accuracy, represented by coefficient of variation (CV) across replicates and the fold change (FC) bias between different loading amounts samples, respectively. Table 1 presents the comparative results of Beta-DIA and DIA-NN on the test datasets, using the same predicted spectral libraries and each software’s default or optimal parameters. Overall, Beta-DIA significantly increases the number of peptide/protein identifications while maintaining comparable data completeness to DIA-NN, particularly on low detection datasets. It also demonstrates superior quantification accuracy, though the quantitative CV and runtime show mixed results between the two tools.

**Tab. 1.** The performance comparison table between Beta-DIA and DIA-NN on test datasets. , All datasets consist of triplicate analyses under different load amounts except for the single-cell datasets Extra-3-1/2. Identification-Gain represents the average increase in the number of identifications across all .d files, achieved by Beta-DIA compared to DIA-NN. Identification-Completeness refers to the average completeness across all triplicates in the dataset. Quantification-CV represents the mean coefficient of variation (CV) across all replicates. Quantification-FC Bias indicates the median of relative deviation between the measured and theoretical FC of the highest and other load amounts (If the maximum loading amount is 200 ng and another sample has a loading amount of 100 ng, the theoretical FC would be 2. If the measured FC is 1.5, the relative deviation of the FC is calculated as (1.5−2)/2=−25%). Time reflects the average analysis time per .d file. Quantification metrics are not applicable for Extra-3-1/2. “X vs. Y” always means DIA-NN vs. Beta-DIA.

| Dataset | Description | Level | Identification |  | Quantification |  | Time |
| --- | --- | --- | --- | --- | --- | --- | --- |
|  |  |  | Gain | Completeness | Precision-CV | Accuracy-FC Bias |  |
| Base-1 | diaPASEF by standard cycle | Pr. | 13.2% | 0.80 vs. 0.79 | 0.12 vs. 0.13 | -24.4% vs. -6.4% | 15.6 vs. 20.8 |
|  |  | Pg. | 4.2% | 0.91 vs. 0.91 | 0.15 vs. 0.13 | -24.6% vs. -8.0% |  |
| Base-2 | diaPASEF by 25% duty cycle | Pr. | 3.5% | 0.83 vs. 0.80 | 0.13 vs. 0.13 | -19.3% vs. -3.0% | 44.9 vs. 30.6 |
|  |  | Pg. | 0.5% | 0.92 vs. 0.91 | 0.16 vs. 0.14 | -20.7% vs. -9.6% |  |
| Base-3 | High throughput | Pr. | 4.6% | 0.78 vs. 0.77 | 0.11 vs. 0.15 | 103.5% vs. 81.7% | 6.5 vs. 14.4 |
|  |  | Pg. | 0.7% | 0.90 vs. 0.89 | 0.17 vs. 0.15 | 107.2% vs. 100.5% |  |
| Base-4 | Dilution series | Pr. | 19.8% | 0.76 vs. 0.77 | 0.11 vs. 0.15 | -48.6% vs. -46.4% | 13.5 vs. 20.4 |
|  |  | Pg. | 7.0% | 0.87 vs. 0.87 | 0.16 vs. 0.14 | -51.3% vs. -51.9% |  |
| Base-5 | Two-proteome experiment | Pr. | 8.6% | 0.84 vs. 0.83 | 0.11 vs. 0.12 | -5.1% vs. -4.3% | 46.9 vs. 54.0 |
|  |  | Pg. | 2.4% | 0.94 vs. 0.94 | 0.15 vs. 0.11 | -6.3% vs. -5.6% |  |
| Extra-1-1 | 3min gradient + dilution | Pr. | 10.9% | 0.72 vs. 0.73 | 0.11 vs. 0.14 | -70.1% vs. -56.8% | 8.8 vs. 11.9 |
|  |  | Pg. | 1.1% | 0.84 vs. 0.84 | 0.18 vs. 0.16 | -74.0% vs. -66.2% |  |
| Extra-1-2 | Two-proteome test (5min gradient) | Pr. | 4.9% | 0.77 vs. 0.78 | 0.12 vs. 0.16 | -9.3% vs. -5.6% | 9.5 vs. 15.5 |
|  |  | Pg. | 1.7% | 0.89 vs. 0.89 | 0.18 vs. 0.16 | -9.6% vs. -5.7% |  |
| Extra-2 | Plasma by timsTOF HT | Pr. | 14.5% | 0.83 vs. 0.81 | 0.14 vs. 0.19 | -34.8% vs. -24.7% | 45.0 vs. 27.2 |
|  |  | Pg. | 13.0% | 0.85 vs. 0.83 | 0.19 vs. 0.18 | -34.7% vs. -28.6% |  |
| Extra-3-1 | Single cell by timsTOF SCP | Pr. | 50.4% | 0.30 vs. 0.30 | — | — | 37.0 vs. 13.1 |
|  |  | Pg. | 32.5% | 0.39 vs. 0.39 | — | — |  |
| Extra-3-2 | Single cell by timsTOF Ultra | Pr. | 47.8% | 0.26 vs. 0.26 | — | — | 27.8 vs. 15.0 |
|  |  | Pg. | 29.0% | 0.36 vs. 0.37 | — | — |  |

Specifically, for the Base-1 dataset (standard cycle, Supplementary Figure 7), Beta-DIA outperformed DIA-NN at 1% FDR, with a 8%-18% increase in peptide identifications and a 2%-8% increase in protein group identifications, while maintaining similar data completeness across three replicates. Both Beta-DIA and DIA-NN achieved median peptide/protein quantification CVs below 20%. In terms of FC between low (10ng and 50ng) and high (100ng) loading amount samples, Beta-DIA’s quantification ratio median are much closer to the expected ratio values compared to DIA-NN, confirming its qualitative accuracy from a quantitative perspective. To further validate the reliability of the gains, we conducted an external FDR calculation using a two-species (human-*A. thaliana*) spectral library on the runs with the minimum and maximum gains achieved by Beta-DIA. We found that, at the 1% external FDR (external FDR = #*A. thaliana* / #human), Beta-DIA achieved a 24-33% gain in peptide identifications and a 3-10% gain in protein group detections; at the same reported FDR, DIA-NN exhibited a smaller external FDR than Beta-DIA. This suggests that, on the Base-1 dataset, Beta-DIA provides a more reliable confidence ranking for target peptides than DIA-NN, though DIA-NN reports a more conservative FDR. Similar results were observed across the Base-2 (25% duty cycle, Supplementary Figure 8), Base-3 (high throughput, Supplementary Figure 9), Base-4 (dilution series, Supplementary Figure 10) and Extra-1 (ultra-high throughput, Supplementary Figure 12) datasets. For Base-4, further Venn diagram analysis (Supplementary Figure 10.c) of the 0.2ng sample revealed that additional protein group identified by Beta-DIA had a higher overlap with those from DIA-NN at 200ng, supporting the reliability of the observed increase. For the known mixing ratio quantitative datasets Base-5 (human + yeast, Supplementary Figure 11) and Extra-1-2 (human + *E. coli*, high throughput, Supplementary Figure 13) datasets, Beta-DIA achieved a modest increase in identifications while demonstrating superior quantitative accuracy. For the Extra-2 (plasma on timsTOF HT, Supplementary Figure 14) dataset, Beta-DIA not only achieved a significant increase, but also outperformed DIA-NN in both quantitative accuracy and FDR control. For the single-cell datasets Extra-3-1 (on timsTOF SCP, Supplementary Figure 15) and Extra-3-2 (on timsTOF Ultra, Supplementary Figure 16), Beta-DIA identified approximately 50% more peptides and 30% more protein groups, with similar FDR control.

In summary, the extensive test data, encompassing various diaPASEF methods, sample types, loading amounts, mass spectrometry instruments, and throughput levels, demonstrate that Beta-DIA comprehensively and reliably outperforms DIA-NN, achieving superior qualitative and quantitative performance for single-shot diaPASEF data.

## 3 Discussion

In this work, we present Beta-DIA for the qualitative and quantitative analysis of diaPASEF data. Beta-DIA achieves significant gains in identification performance by applying learning-based scoring to complex features such as 2D coelution consistency and spectrum similarity, in combination with other function-based scoring methods. Beta-DIA integrates deep learning into the core of the analysis software—the scoring, representing another application of deep learning beyond spectral library prediction. Moreover, this integration does not lead to a significant increase in analysis time. Meanwhile, Beta-DIA further enhances the quantitative accuracy of single-shot diaPASEF data.

Our results also indicate that the iterative development of the timsTOF platform has not affected the usability of Beta-DIA, which is based on a pretrained model. We also found that using heterogeneous large-scale data may outperform using experiment-specific data to pretrain DeepProfile (see Methods and Supplementary Figure 6). Given that Beta-DIA includes a refine process, we believe for most cases Beta-DIA eliminates the need for users to train models with localized data, enabling the software to be ready to use out-of-the-box.

A natural question that arises is whether the pretrained model-based workflow can be extended to DIA data analysis on other mass spectrometry platforms. If feasible, this will demonstrate the generalizability of the pretrained model across different mass spectrometry platforms and varying data complexities. If not, an alternative could be the on-the-fly model training approach, where function-based scoring completes the initial identification, and the results are then used to train a learning-based scoring model for the final hybrid scoring and identification, as exemplified by AlphaXIC.

Furthermore, the concept of using deep learning to score spectrum similarity in this work can clearly be extended to any scenario requiring spectrum similarity calculations, such as in DDA identification or mass spectrometry-based metabolomics.

In summary, it is anticipated that integrating learning-based and function-based scoring will yield deeper identification results than function-based scoring alone, thus providing maximal support for downstream proteomics applications.

## 4 Methods

### Identification algorithms

Beta-DIA takes as input a spectral library and .d files. Currently, Beta-DIA supports directly reading spectral libraries in the .speclib or .parquet format as defined by DIA-NN, from which it retrieves target peptide ions and its fragment ion information (including RT, ion mobility, *m/z*, and fragment ion intensities). Beta-DIA utilizes AlphaTims^[23]^ to read .d files as raw profile data, and converts the profile data into centroided data according to DIA-NN’s 2D peak-picking algorithm.

The identification process of Beta-DIA can be divided into four stages: the calibration stage, the first round of identification, the second round of identification and the polish of results.

In the calibration stage, Beta-DIA extracts chromatographic elution groups for each target peptide ion (including the precursor in MS1 scan, the unfragmented precursor and the top 12 fragment ions in MS2 scan, 14 ions in total) from the centroided diaPASEF data with a fixed width in *m/z* (20 ppm) and ion mobility (0.05 Vs cm^-2^) across the whole RT scale. Beta-DIA then calculates the coelution consistency of elution groups using cosine function within a sliding window, with a window width of 13 cycles and a sliding step of 1 cycle. Beta-DIA selects the elution time, ion *m/z*, and ion mobility values corresponding to the apex of the elution group with the highest coelution consistency score as the measured attributes of the target precursors. Based on the measured and the spectral library attributes of target precursors, Beta-DIA constructs RT, *m/z*, and mobility error profiles and uses the Calib-RT algorithm^[24]^, a local regression (LOESS) algorithm with noise removal, to calibrate them. It is important to note that FDR control is absent during this calibration stage, which relies on a “rough” identification result that includes false positives.

Next is the first round of identification. For each target peptide in the spectral library, Beta-DIA generates a decoy peptide via the mutate method (GAVLIFMPWSCTYHKRQEND to LLLVVLLLLTSSSSLLNDQE mutation pattern is used). For each target peptide or decoy peptide, Beta-DIA extracts chromatograms of 14 monoisotopic ions from the centroided diaPASEF data with a fixed width in RT (1/15 of total gradient), *m/z* (20 ppm) and ion mobility (0.05 Vs cm^-2^). Beta-DIA then seeks potential elution groups and assigns function-based scores to them. Meanwhile, Beta-DIA uses a pretrained DeepProfile model to perform 2D (RT and mobility) coelution consistency scoring on 2D elution groups which are retrieved from profile data. Specifically, DeepProfile includes two models: DeepProfile-14 and DeepProfile-56 (unless otherwise specified, DeepProfile refers to both DeepProfile-14 and DeepProfile-56). DeepProfile-14 scores the 2D elution groups of the 14 ions mentioned earlier under −1, 0, +1, and +2 neutron conditions individually, while DeepProfile-56 scores the 2D elution groups of the 14*4 ions as a whole. After scoring, Beta-DIA uses an ensemble neural network to distinguish between target and decoy peptides, resulting in a single discrimination score for each peptide, by which the FDR is calculated (#decoy/#target). The ensemble neural network consists of twelve neural network-based binary classifiers, each treating target peptides as positive samples and decoy peptides as negative samples during training. Each neural network comprises a series of hidden layers (consistent with DIA-NN using ^[25, 20, 15, 10, 5]^ neurons) and optimizes parameters by the Adam optimizer (with a learning rate of 0.001). Importantly, the dataset is split into a training set and a validation set in a 4:1 ratio. The primary difference between each classifier lies in the variation of training batch sampling (batch size 50 by default). Training stops when the accuracy on the validation set does not improve for five consecutive epochs (*i.e.*, train patient = 5) or reaches a maximum of 10 training epochs.

Then is the second round of identification. Beta-DIA constructs local training data based on the results of the first round of identification to refine DeepProfile (scoring 2D coelution consistency) and train DeepMall (scoring spectrum similarity) from scratch. Specifically, the best elution groups of target precursors at 1% FDR from the first round are labelled as positive samples, and the second-best elution groups are labelled as negative samples, both of which are used to construct the corresponding 2D elution groups and spectrum similarity data. Based on these samples data, DeepProfile is refined and DeepMall is trained. The scores from DeepProfile and DeepMall are then integrated with other scores from the first round into an ensemble neural network, and the FDR is updated. Both the refinement of DeepProfile and the training of DeepMall use the Adam optimizer (learning rate of 0.0001). The samples are split into a training set and a validation sets in a 4:1 ratio. The batch size is 64. Training is halted when the accuracy on the validation set does not improve for 5 consecutive epochs or after a maximum of 50 training epochs. Notably, the refinement of DeepProfile involves only the layers which are used for calculating class probabilities, while the other feature layers remain frozen to preserve the feature extraction capabilities learned during the pretraining phase. It is important to note that neither the refinement of DeepProfile nor the training of DeepMall uses decoy information, thus eliminating the possibility of decoy data leakage compromising FDR accuracy. We supplemented ablation experiments comparing learning-based scoring and function-based scoring (see Supplementary Figure 3), which demonstrate that: a) DeepProfile-14 and DeepProfile-56 are complementary rather than hierarchical; b) the learning-based scoring by DeepProfile and DeepMall is complementary to function-based scoring; and c) the pretraining and refinement of DeepProfile are mutually complementary.

The final stage involves polishing the results from the second round of identification to reduce the redundant use of fragments for multiple precursors from the same spectrum. Beta-DIA checks whether the fragment ions of each target precursor could belong to a target precursor with a higher discriminative score. When two target precursors are located within the same fragmentation window, and their RT deviation, ion mobility deviation, and *m/z* deviation of corresponding fragment ions fall within predefined ranges, the fragment ion of the target precursor with the lower discriminative score is considered suspicious. If a target precursor contains at least three suspicious fragment ions, or if the total coelution score of suspicious fragment ions exceeds two-thirds of the total coelution score of all fragment ions, the target precursor is discarded. Then, the ensemble neural network updates the calculation of FDR based on the remaining target and decoy precursors.

Protein identification begins with the IDPicker^[25]^ algorithm to assign peptides to protein groups (*i.e.*, when a peptide is associated with multiple proteins, it is preferentially assigned to the protein with the most peptides at 5% FDR). Next, a single discrimination score is calculated for each protein group using the formula 1-∏(1-P_i_), where P_i_ is the peptide discrimination score. Finally, the FDR for the protein group is calculated using the #decoy/#target method.

### Quantification algorithms

After Beta-DIA performs the aforementioned identification process, it obtained the best elution group for each peptide ion by the fixed tolerance width in *m/z* (20ppm) and ion mobility (0.05 Vs cm^-2^). Beta-DIA then calculates the correlation between each pair of elution profiles and selects the profile with the highest correlation to others as the best profile. Next, Beta-DIA iterates over different mass and mobility tolerances to re-extract the elution profiles of ions, selecting the elution profile with the highest correlation to the best profile as the final extracted profile. The mass tolerance space is ^[20, 16, 12, 8, 4]^ ppm, and the mobility tolerance space is [0.02, 0.01] Vs cm^-2^ (starting at 0.02 instead of 0.05 because the ion’s measured mobility was obtained during the identification phase). The final quantification of the peptide ion is determined by the sum of the areas under the profiles of the top-6 fragment ions.

For protein group quantification, the confidences of the peptides it contains are taken into account. If a protein group contains peptides within 1% FDR, its quantification is determined by the average of the top-3 peptides at an FDR of 1%; if it does not contain peptides within 1% FDR, the protein group’s quantification is based on its top-1 peptide. Here, top-n refers to ranking peptides in descending order by their quantities.

### DeepProfile

DeepProfile is used to score the 2D coelution consistency on 2D elution groups. The pretraining data (Supplementary Table 3) of DeepProfile are provided by Shanghai Omicsolution Co., Ltd. These sixteen runs used for training are designed to maximize data diversity by combining different sample types, gradients and window slicing. They were identified using DIA-NN, which determined the best elution group for each peptide at 1% FDR. To augment the data, each best elution group was shifted by −1, 0, and 1 cycles, resulting in ∼2.5M 2D elution groups as positive samples. We then picked the top-3 suboptimal elution group based on coelution scoring using cosine similarity across the whole elution time as negative samples. In this way, the number ratio of positive to negative samples keeps 1:1. The 2D elution groups with dimensions of 14×13×50 (where 14 refers to the aforementioned ions, 13 to the cycles, and 50 to the bins of 0.001 Vs cm^-2^ width) were used to create the training dataset for DeepProfile-14. When accounting for isotopic variations, the 14 ions expand to 56 ions, and the corresponding 2D elution groups with dimensions of 56×13×50 form the training dataset for DeepProfile-56.

Although DeepProfile-14 and DeepProfile-56 have different input dimensions, their model designs are identical (Supplementary Figure 1). The raw 2D elution groups are first normalized at both the ion level and the overall level. The normalized data then pass through two rounds of [convolution, max pooling] and a fully connected layer, producing a 16-dimensional feature vector. Two 16-dimensional feature vectors are concatenated with an embedding representing the actual number of fragment ions, resulting in a 48-dimensional vector. This vector undergoes two fully connected transformations to produce the final class probability values. During the refinement process for DeepProfile, the two 16-dimensional feature vector is frozen and remains unchanged.

DeepProfile’s training is implemented by PyTorch, employing the Adam optimizer (with an initial learning rate of 0.001 and 1024 samples per batch) and a cross-entropy loss function. The training data is split into training and validation sets in a 4:1 ratio. Training stops when the validation accuracy does not improve for 5 consecutive epochs or after a maximum of 50 training epochs. Supplementary Figure 4 illustrates the impact of training set size on model performance, with the validation set size kept constant. It shows that when the training set reaches 40% of its maximum size, adding more training data does not result in higher validation accuracy, suggesting that the current data size is sufficient for effective model training. Supplementary Figure 5 shows the impact of different training epochs on identification results. It can be observed that additional training epochs (e.g., 50 epochs) do not lead to the identification of more peptides, indicating that the early stopping method based on a patience of 5 epochs is reasonable.

### DeepMall

Compared to coelution consistency, spectrum similarity is a secondary feature in the identification process. Therefore, DeepMall (Supplementary Figure 2), designed to score spectrum similiarity, is not pre-trained but rather trained on-the-fly. After determining the best elution group for each peptide at 1% FDR, Beta-DIA constructs the input data for DeepMall based on the top-12 fragment ions, including: a) the relative intensities of the fragment ions provided by the spectrum library; b) fragment ions type (b ions coded as 1, y ions coded as 2); c) normalized measurements intensities of the fragment ions from the three cycles around the elution apex, as well as the corresponding mass errors (normalized by dividing by 20 ppm) and mobility errors (normalized by dividing by 0.05 Vs cm^-2^); d) cosine similarity scores of elution profiles; e) signal-to-noise ratios of elution profiles; f) profile areas. DeepMall treats these 14 features for each fragment ion as a whole and feeds them into a two-layer bidirectional GRU, which ultimately outputs a probability value indicating spectrum similarity. Since the spectrum similarity calculation incorporates not only the spectrum library intensities and measured intensities but also as much additional information about the fragment ions as possible to provide a confidence weight for spectrum similarity, the model is named DeepMall, reflecting its comprehensive amalgamation of ion information. The description of DeepMall’s training can be found in the “Identification algorithms” section.

### Pretraining DeepProfile with experiment-specific datasets

We downloaded a urine cohort experimental dataset^[26]^, which contains 37 .d files, each identified using DIA-NN. Except for the two files with the minimum and maximum identifications by DIA-NN, 35 files were processed to generate experiment-specific pretraining data using the aforementioned method, resulting in ∼3M 2D elution groups as positive samples. We applied the same pretraining method to complete the pretraining of the DeepProfile model based on experiment-specific data. Then, Beta-DIA analyzed the two excluded files using DeepProfile pretrained on the heterogeneous data or the experiment-specific data, with results shown in Supplementary Figure 6. As can be seen, the DeepProfile model pretrained on heterogeneous data slightly outperforms the one trained on experiment-specific data. Given that Beta-DIA includes a refine process, it can be considered that, to some extent, Beta-DIA eliminates the need for users to pretrain DeepProfile using experiment-specific datasets.

### Spectral library generation

For all analyses, we used DIA-NN to theoretically digest protein sequences and generate predicted spectral libraries. *Homo sapiens* (taxon identifier: 9606, 20,420 sequences), *Mus musculus* (taxon identifier: 10090, 17,212 sequences), *S. cerevisiae* (taxon identifier: 4932, 6,060 sequences), *E. coli* (taxon identifier: 562, 4,401 sequences) and *A. thaliana* (taxon identifier: 3702, 16,310 sequences) from UniProt (reviewed sequences only; downloaded on June, 2024) were used. The enzymatic digestion and prediction parameters by DIA-NN were as follows which are self-explanatory: ‘--min-fr-mz 200 --max-fr-mz 1800 --met-excision --min-pep-len 7 --max-pep-len 30 --min-pr-mz 300 --max-pr-mz 1800 --min-pr-charge 1 --max-pr-charge 4 --cut K*,R* --missed-cleavages 1 --unimod4 --var-mods 1 --var-mod UniMod:35,15.994915,M’. As a result, the predicted spectral library generated for Base-1/2/3/4, Extra-1-1, Extra-2 and Extra-3 contained 5,628,093 precursors; for the two-proteome Base-5 (HeLa + Yeast) contained 7,123,532 precursors; and for the two-proteome Extra-1-2 (K562 + *E. coli*) contained 6,322,592 precursors. The spectral library used for external FDR validation (*Homo sapiens* + *A. thaliana*) contained 9,557,086 precursors.

### Software versions and parameters

DIA-NN with version 1.9.2 was used to predict spectral libraries. Both DIA-NN (version 1.9.2) and Beta-DIA (version 0.7) analyzed each run individually using their respective default parameters.

### Data analysis

Beta-DIA and DIA-NN each generated a report table for every diaPASEF data, with consistent column names across both. Identified peptides and protein groups were filtered at 1% FDR using ‘Q.Value’ and ‘PG.Q.Value’, respectively. Triplicate results were merged using the “at least two” (where a peptide or protein group must exist in at least two of the three replicates) and the average intensity methods.

### Runtime comparison

The runtimes for Beta-DIA and DIA-NN were measured on a Linux desktop equipped with an Intel Core i9-14900KF CPU (32 logical cores), 256 GB of RAM, and an NVIDIA GeForce RTX 4090 GPU. Beta-DIA was set to use 12 CPU cores, while DIA-NN was allocated 24 CPU cores.

### Code availability

Beta-DIA is available for download from https://github.com/YuAirLab/beta_dia and can be installed from PyPi. Python scripts for result summarization and figure generation can be found at the same repository.

## Acknowledgements

We thank Liu Zhu and Catherine C. L. Wong for their early support of this project. This work was funded by National Natural Science Foundation of China [T2222007 to X.W.] and Outstanding Postdoctoral Program of Jiangsu Province, China.

## Supplementary Information

**Supplementary Tab. 1.** Table of detailed descriptions of test datasets.

| Dataset | ID | Description | Sample | Load amount (ng) | Gradient/Throughput | Device (timsTOF) | #Runs |
| --- | --- | --- | --- | --- | --- | --- | --- |
| Base<sup>[1,2]</sup> | 1 | diaPASEF by standard cycle | HeLa | 10-50-100 | 95 min | Pro | 9 |
|  | 2 | diaPASEF by 25% cycle scheme | HeLa | 10-50-100 | 95 min | Pro | 9 |
|  | 3 | High throughput | HeLa | 200 | 60-100-200 SPD | Pro | 9 |
|  | 4 | Dilution series | HeLa | 0.2-1-10-50-200 | 93 min | Pro2 | 15 |
|  | 5 | Two-proteome experiment | HeLa + Yeast | 200 + 45/15 | 90 min | Pro | 6 |
| Extra-1<sup>[3]</sup> | 1 | 3min gradient + dilution | K562 | 10-3000 | 3 min(400 SPD) | Pro | 27 |
|  | 2 | Two-proteome experiment + High throughput | K562 + *E. coli* | 2000 + 500/333 | 5 min(240 SPD) | Pro | 6 |
| Extra-2<sup>[4]</sup> | 1 | Plasma by timsTOF HT | Plasma | 100-300-600-900-1200 | 30 SPD | HT | 15 |
| Extra-3<sup>[5]</sup> | 1 | Single cell by timsTOF SCP | HEK-293T | Single cell | 40 SPD | SCP | 24 |
|  | 2 | Single cell by timsTOF Ultra | HEK-293T | Single cell | 40 SPD | Ultra | 42 |

**Supplementary Tab. 2.**
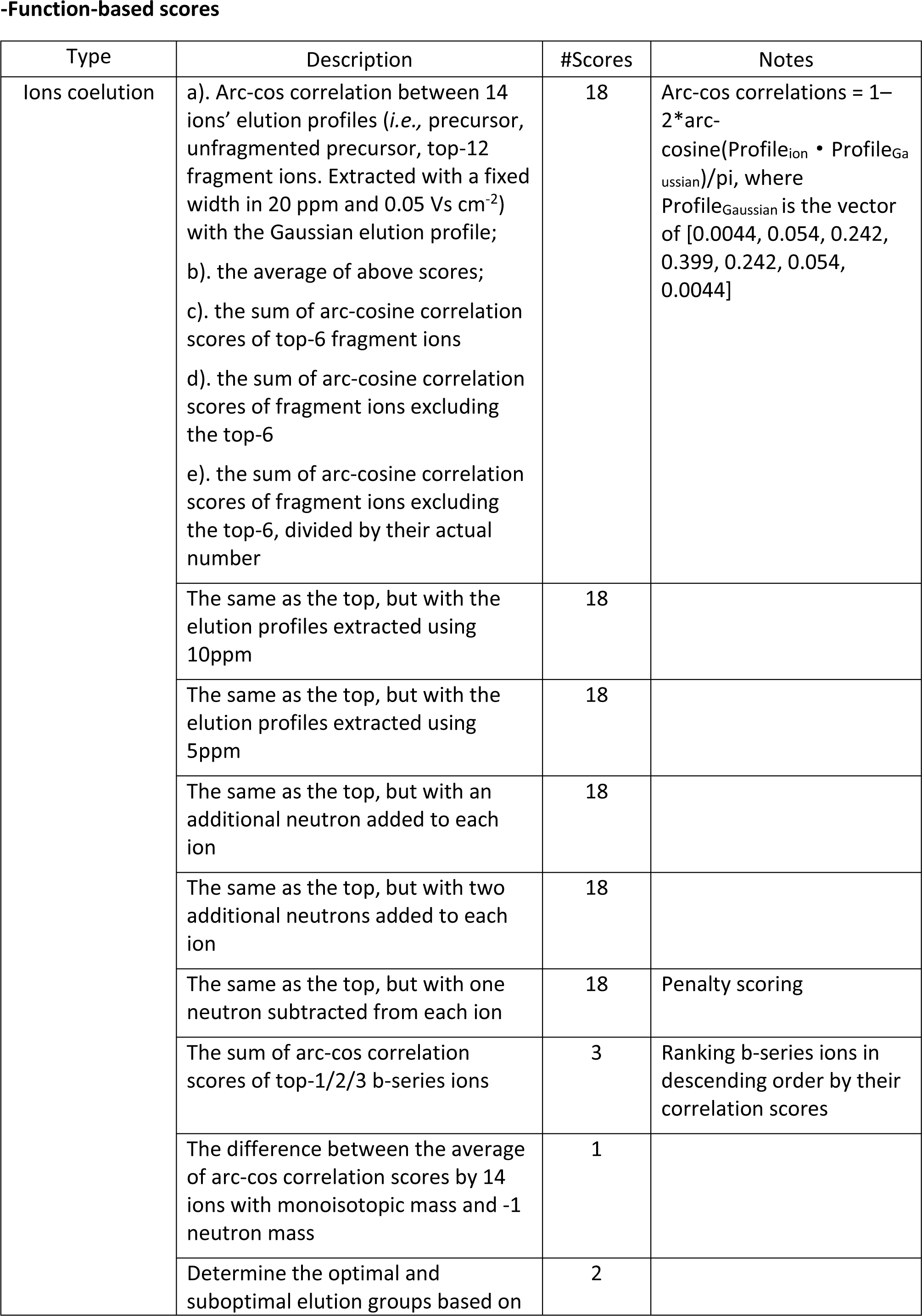
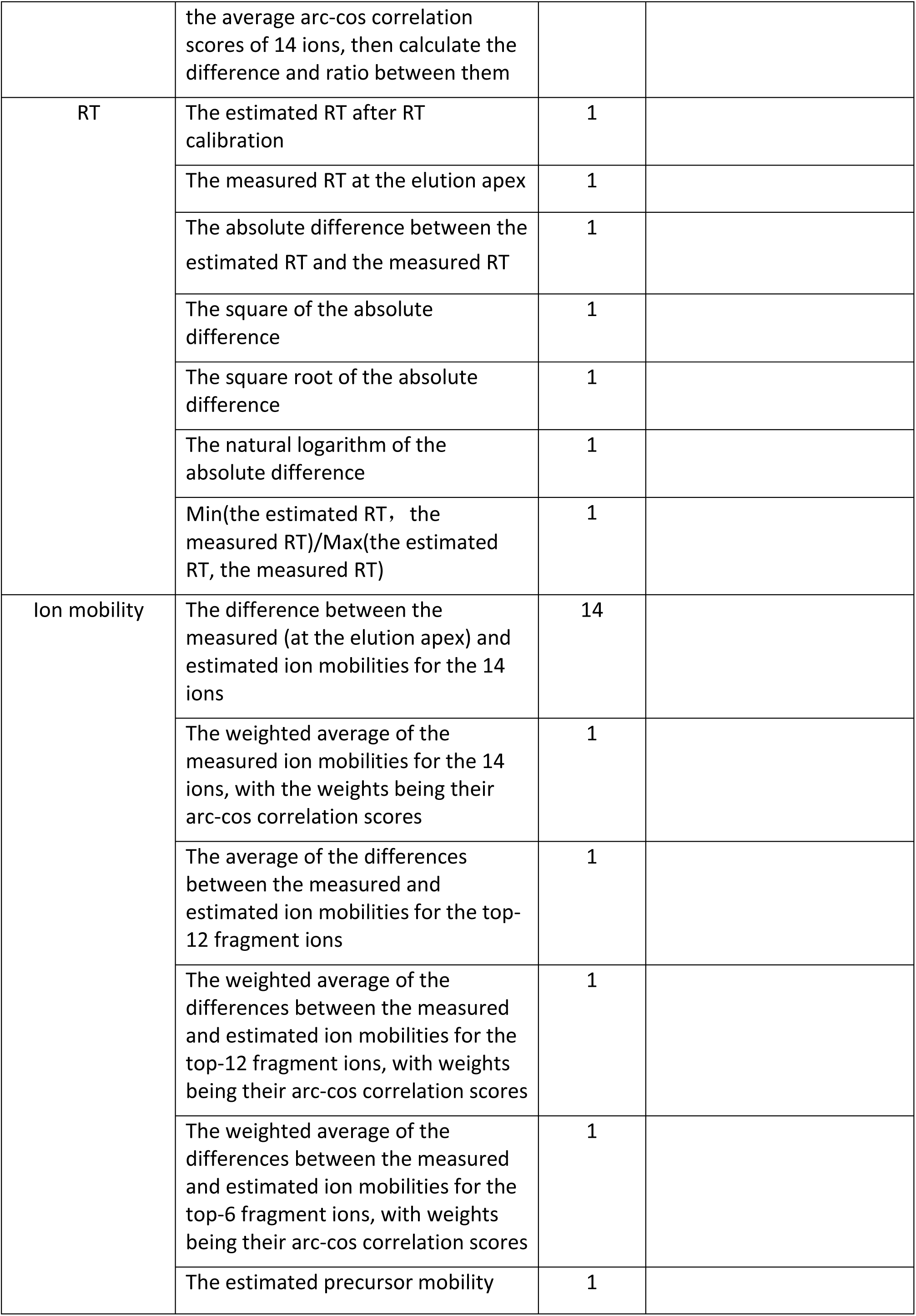
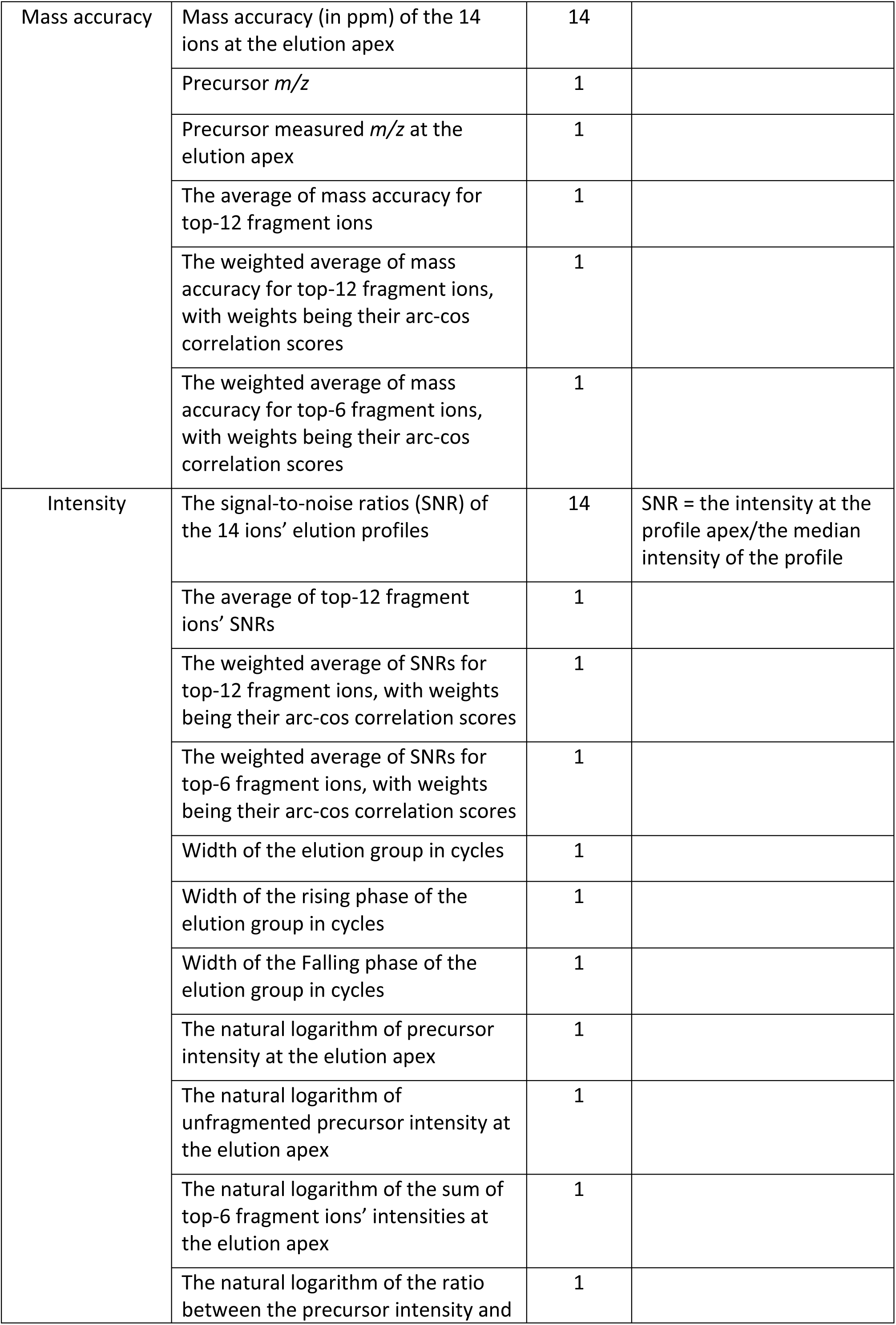
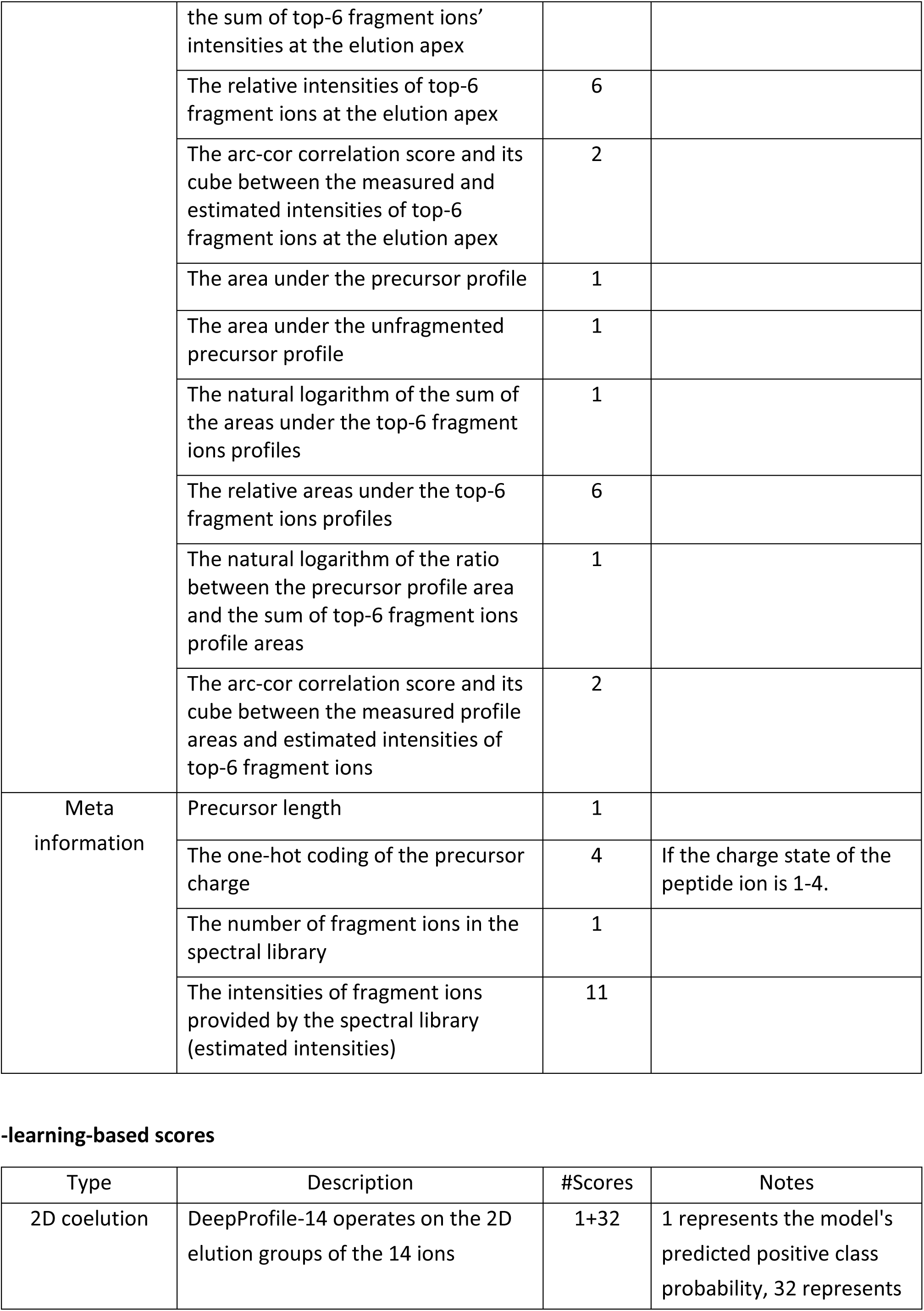
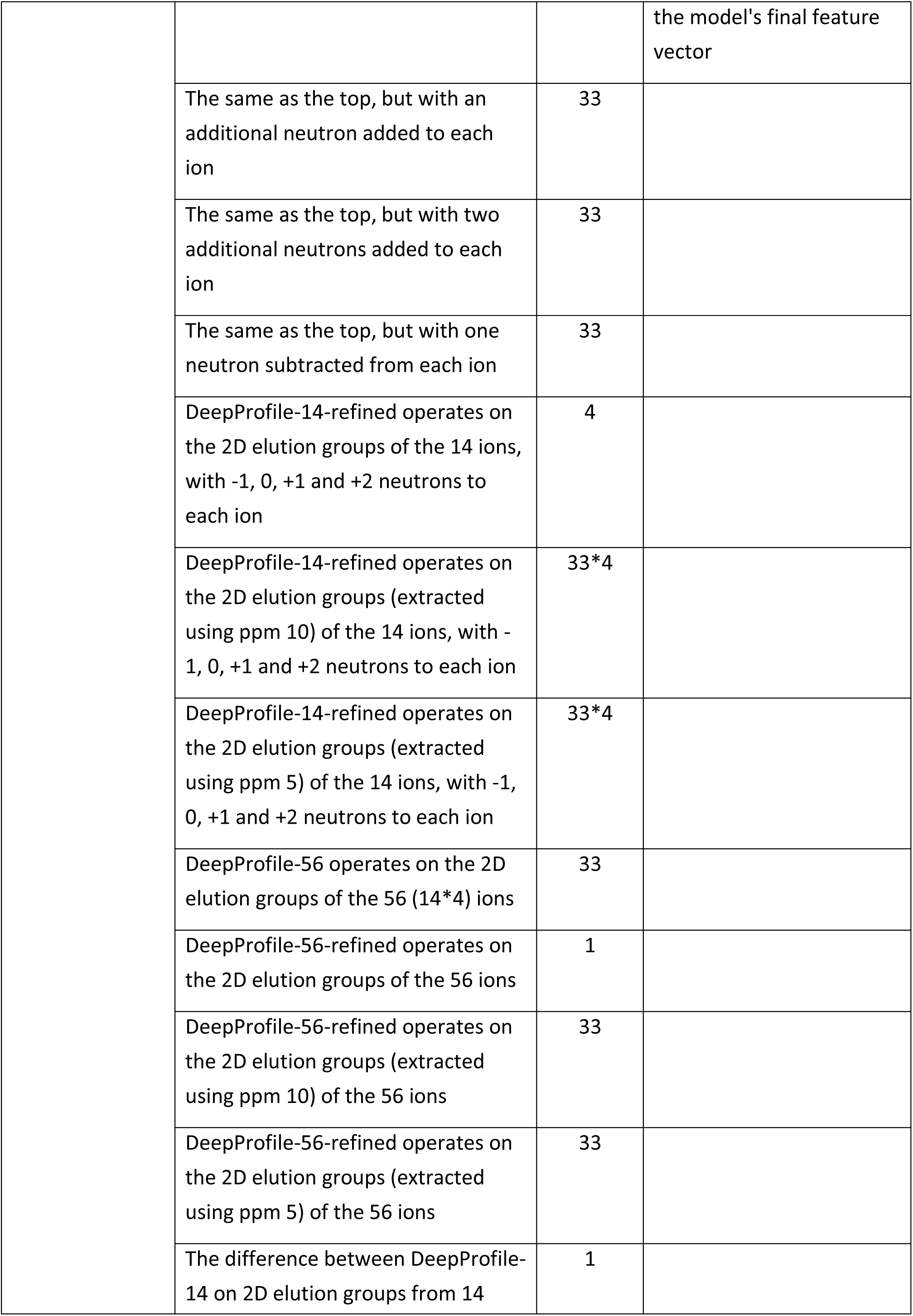
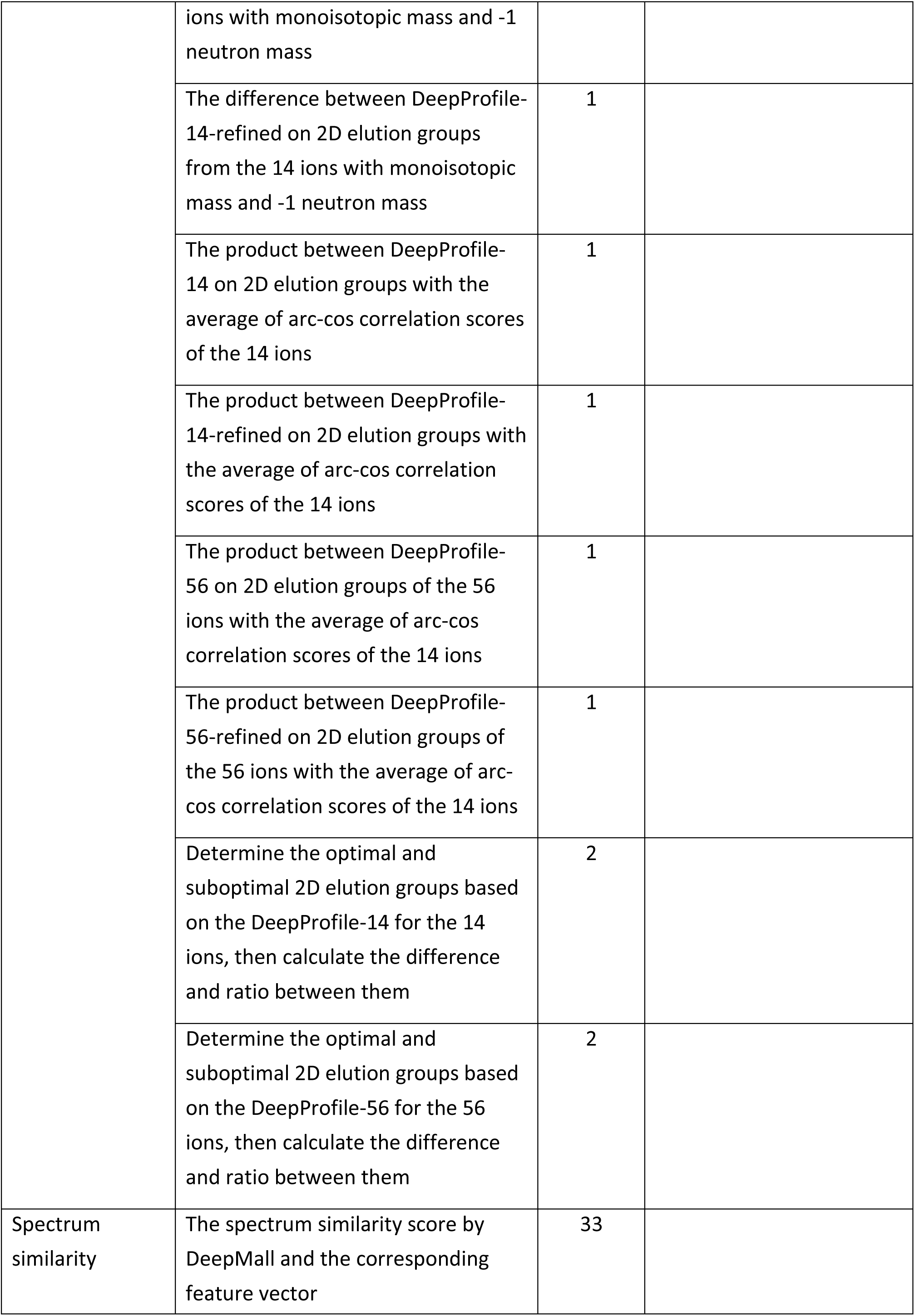
Detailed descriptions of all scoring items.

**Supplementary Tab. 3.**
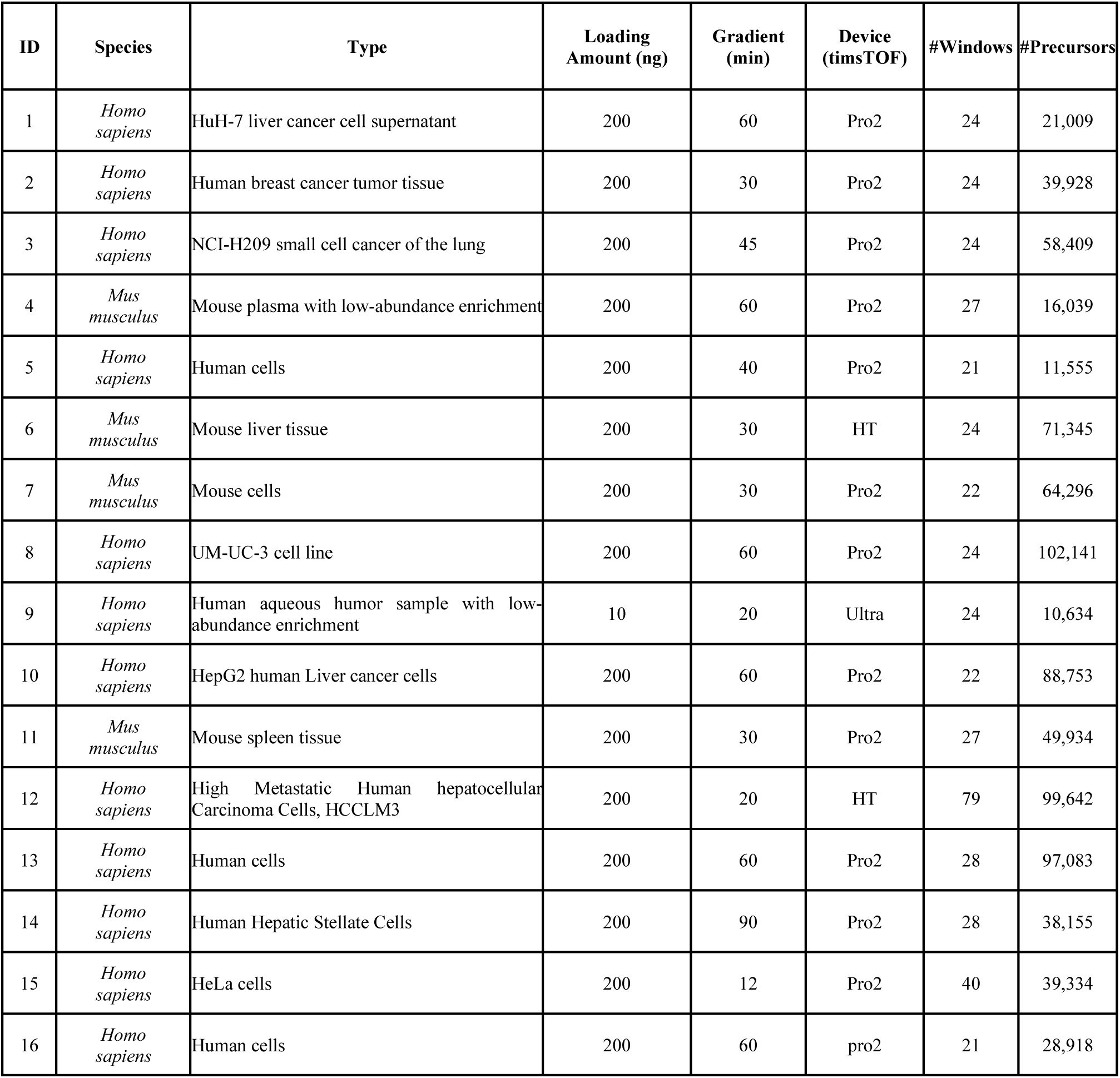
Table of detailed descriptions of training datasets for DeepProfile.

**Supplementary Fig. 1.**
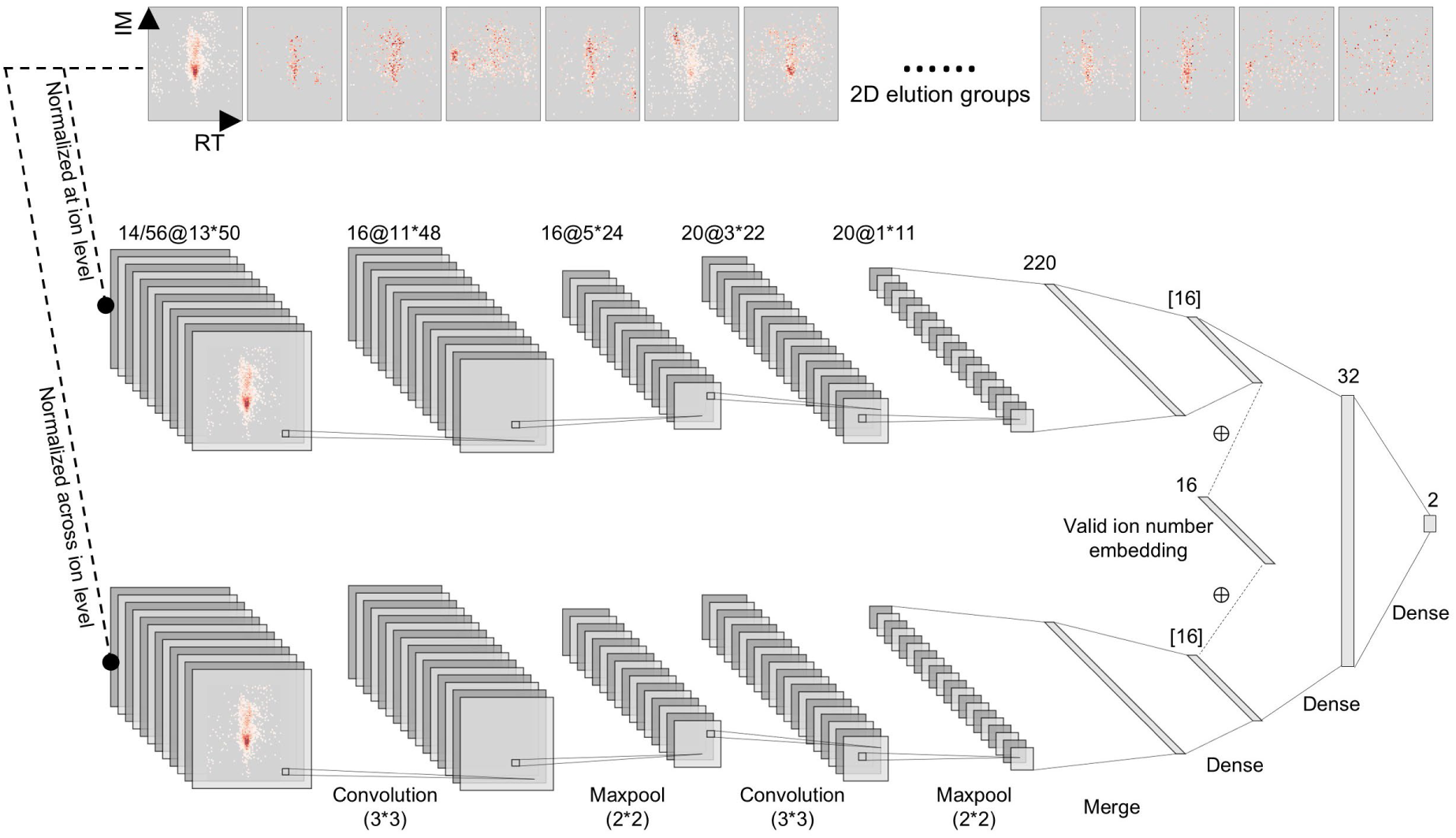
The DeepProfile deep learning model. Deepprofile includes DeepProfile-14 and DeepProfile-56. DeepProfile-14 scores the consistence of 2D elution groups of the 14 ions (precursor, unfragmented precursor, top-12 fragment ions) under −1, 0, +1, and +2 neutron conditions individually. DeepProfile-56 scores the consistency of all these ions’ 2D elution groups as a whole. Each 2D elution group has dimensions of ^[13, 50]^, where 13 refers to the cycles, and 50 to the bins of 0.001 Vs cm^-2^ width. [] indicates feature vectors extracted by the deep learning model, which will be used as part of Beta-DIA’s learning-based scores. The network parameters before feature vectors remain unchanged during the refine stage.

**Supplementary Fig. 2.**
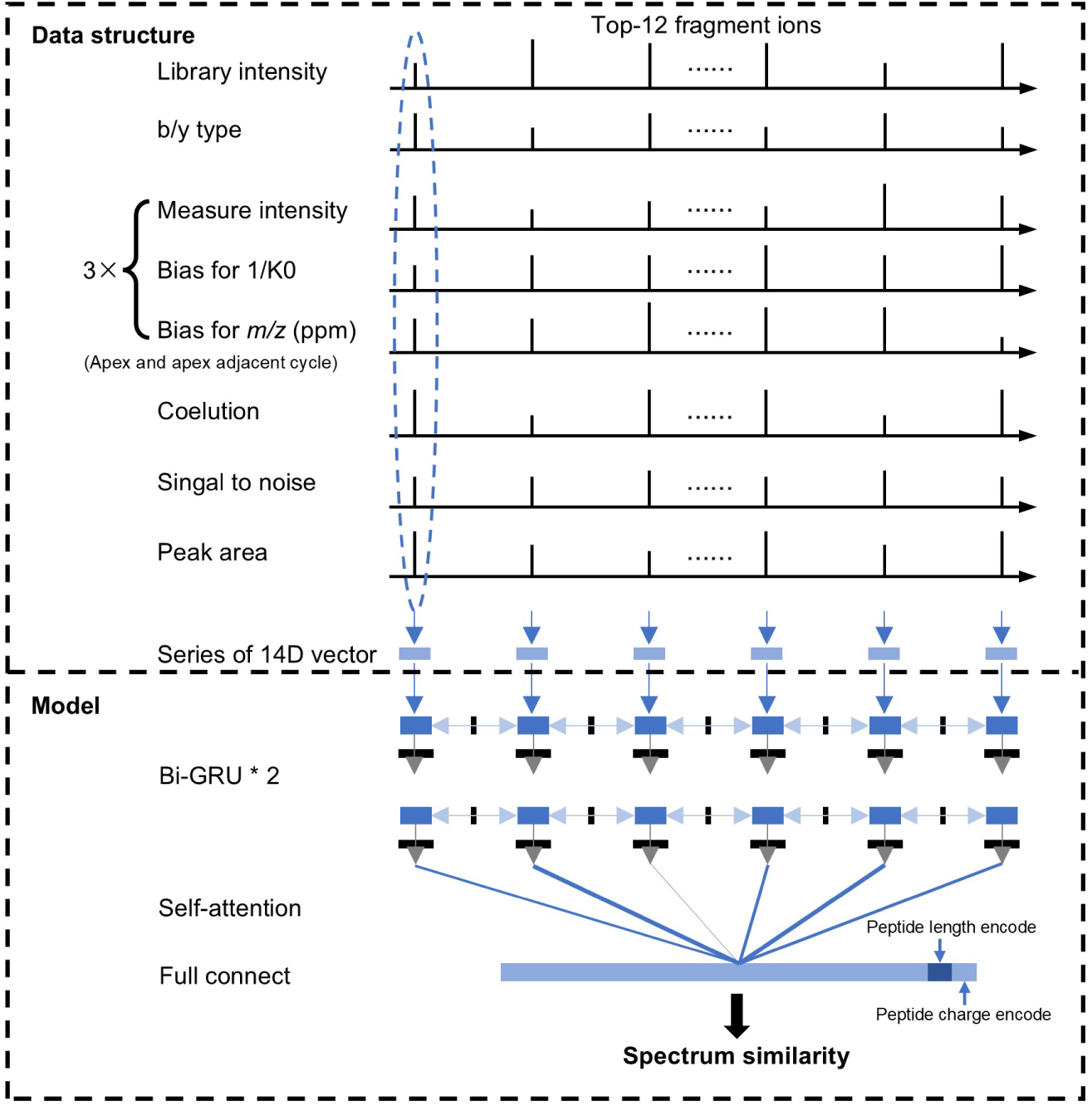
The DeepMall deep learning model. DeepMall is used to score the spectrum similarity based on the top-12 fragment ions. The input of DeepMall includes: a) the relative intensities of the fragment ions provided by the spectrum library; b) fragment ions type (b ions coded as 1, y ions coded as 2); c) normalized measurements intensities of the fragment ions from the three cycles around the elution apex, as well as the corresponding mass errors (normalized by dividing by 20 ppm) and mobility errors (normalized by dividing by 0.05 Vs cm^-2^); d) cosine similarity scores of elution profiles; e) signal-to-noise ratios of elution profiles; f) profile areas. DeepMall treats these 14 features for each fragment ion as a whole and feeds them into a two-layer bidirectional GRU, which ultimately outputs a probability value indicating spectrum similarity.

**Supplementary Fig. 3.**
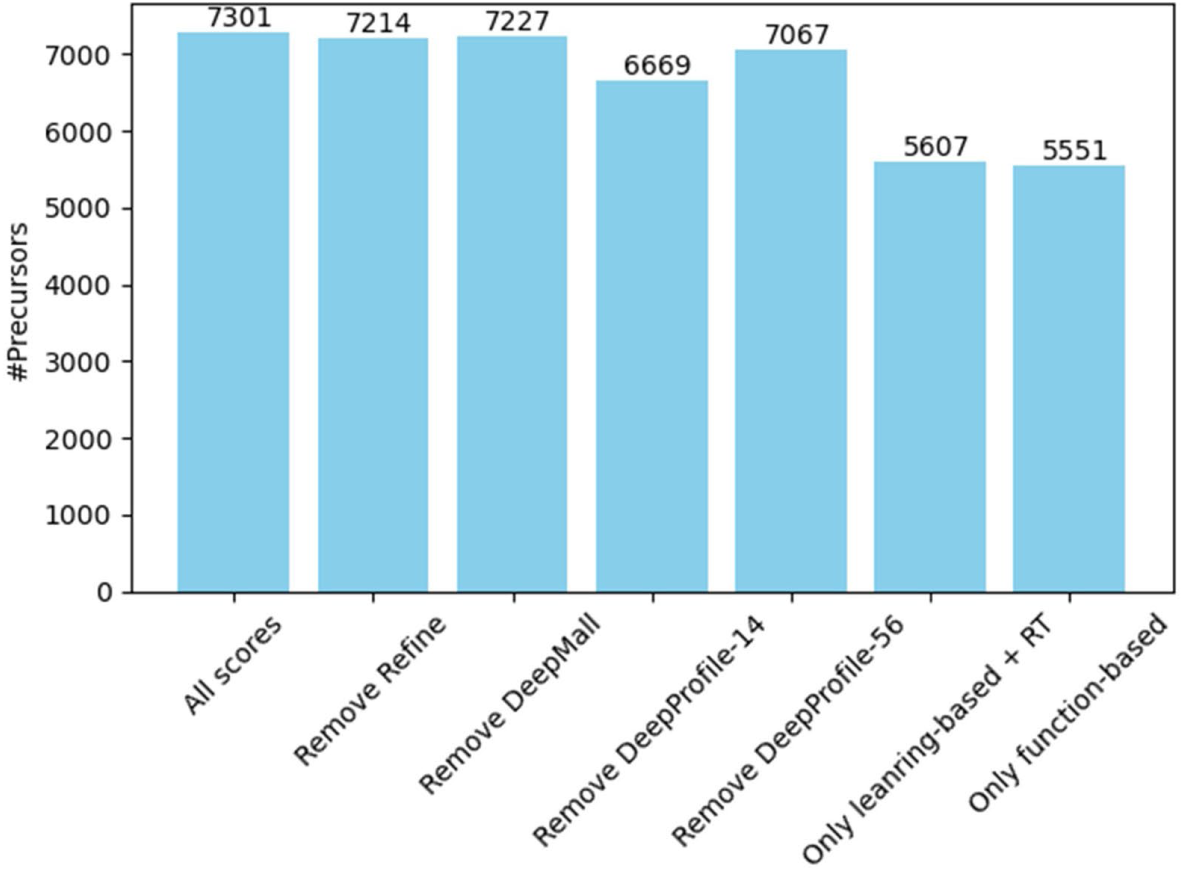
Results of ablation experiments. The bar chart represents the number of peptide identifications at 1% FDR by Beta-DIA under different score combinations on Base-4 dataset (20211101_PRO2_LS_05_MA_HeLaSCS_0.2_ngHS_GE2_1_1409.d). ‘’*All scores*’’ refers to using all function-based and leaning-based scores. ‘’*Remove refine*’’ indicates the removal of DeepProfile-refined related scores. ‘’*Remove DeepMall*’’ indicates the removal of DeepMall related scores. ‘’*Remove DeepProfile-14*’’ indicates the removal of DeepProfile-14 related scores. ‘’*Remove DeepProfile-56*’’ indicates the removal of DeepProfile-56 related scores. ‘’*Only learning-based + RT*’’ refers to using only learning-based and RT-related scores. ‘’*Only function-based*’’ refers to using only functional-based scores. The result demonstrate that: a) DeepProfile-14 and DeepProfile-56 are complementary rather than hierarchical; b) the learning-based scoring by DeepProfile and DeepMall is complementary to function-based scoring; and c) the pretraining and refinement of DeepProfile are mutually complementary.

**Supplementary Fig. 4.**
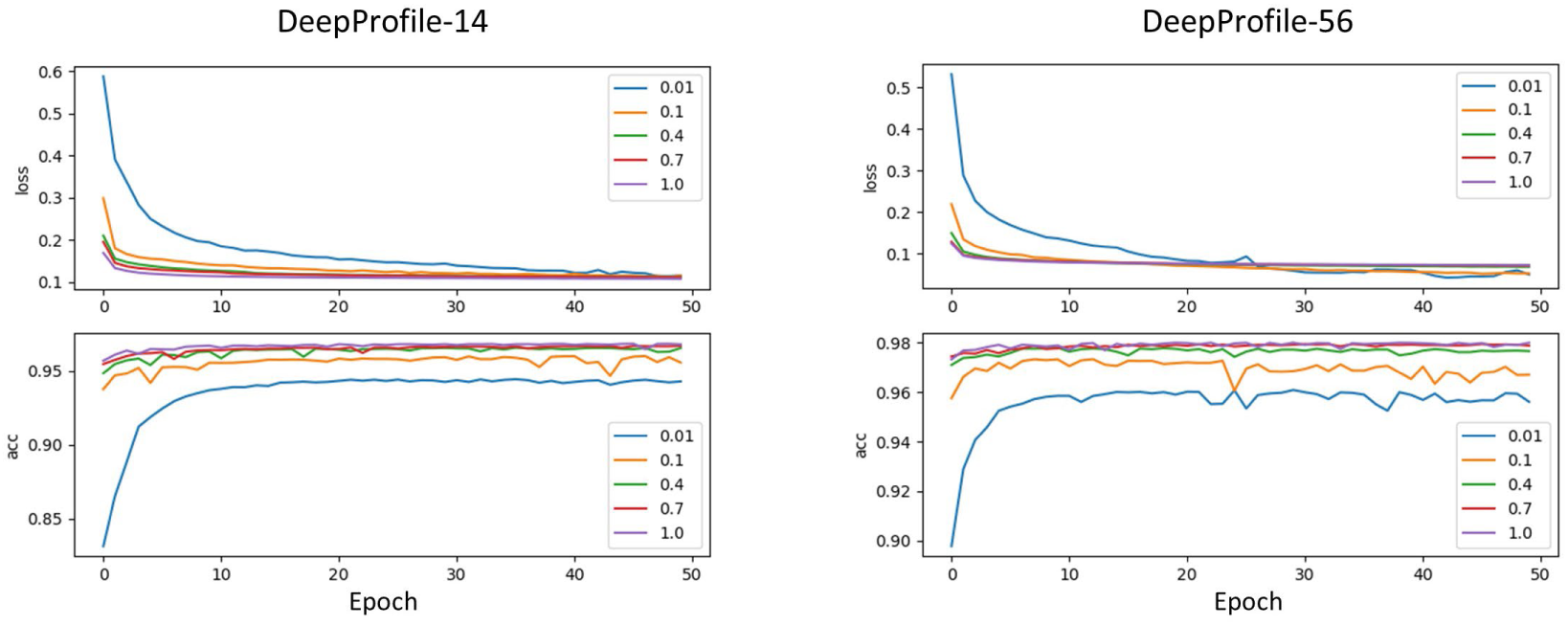
Pretraining results of the DeepProfile under different training data scales and epochs. The upper and lower subplots show the model’s loss on the training set and classification accuracy on the validation set, respectively. The numbers in the legend represent the scale of the training data, where 1 indicates using 100% of the training data, and 0.1 indicates using 10%. Regardless of the training data scale, the validation set remains unchanged. Based on the early stopping criterion with patient-5, DeepProfile-14 ultimately uses the model parameters from epoch 25, while DeepProfile-56 uses the model parameters from epoch 17.

**Supplementary Fig. 5.**
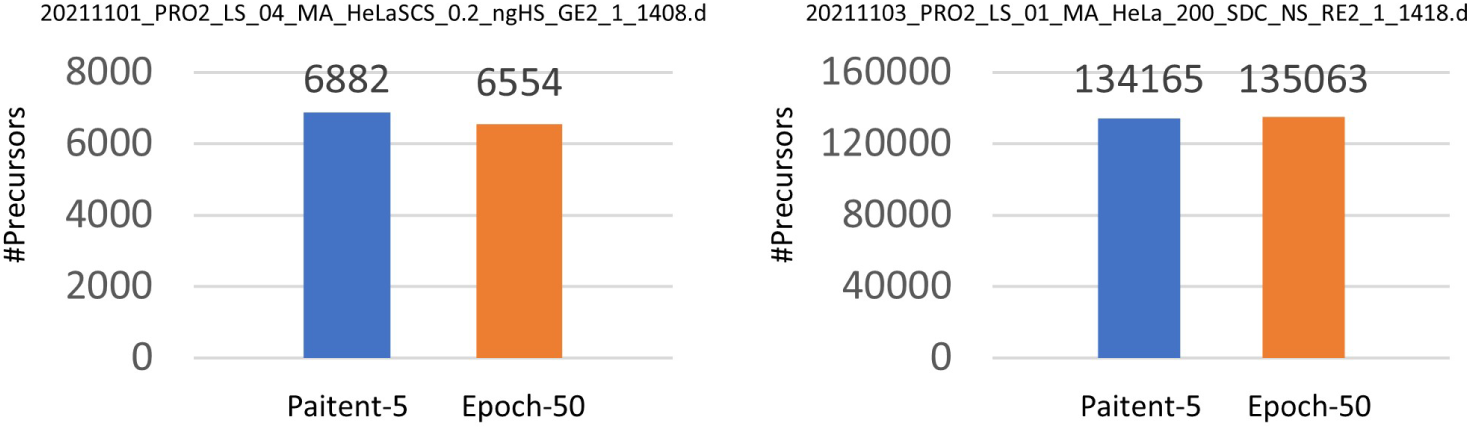
The impact of different training epochs for DeepProfile on identification results. The two .d files from Base-4 dataset, 20211101_PRO2_LS_04_MA_HeLaSCS_0.2_ngHS_GE2_1_1408.d and 20211103_PRO2_LS_01_MA_HeLa_200_SDC_NS_RE2_1_1418.d represent low and high peptide detections, respectively. “Patient-5” refers to using the parameters from epoch 25 for DeepProfile-14 and epoch 17 for DeepProfile-56. “Epoch-50” refers to both models using the parameters from epoch 50.

**Supplementary Fig. 6.**
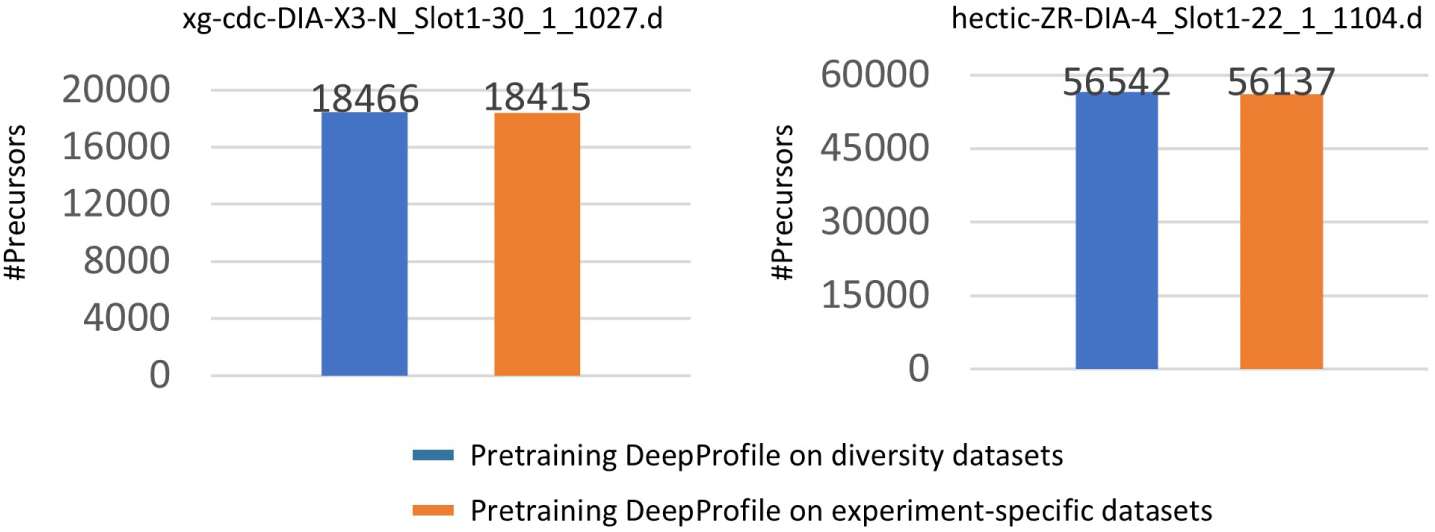
The impact of pretraining DeepProfile using heterogeneous (∼2.5M 2D elution groups) or experiment-specific datasets (∼3M 2D elution groups) on identification results. The two .d files from experiment-specific dataset, xg-cdc-DIA-X3-N_Slot1-30_1_1027.d and hectic-ZR-DIA-4_Slot1-22_1_1104.d represent low and high peptide detections, respectively. Of note, these two .d files are excluded when pretraining DeepProfile using experiment-specific datasets.

**Supplementary Fig. 7.**
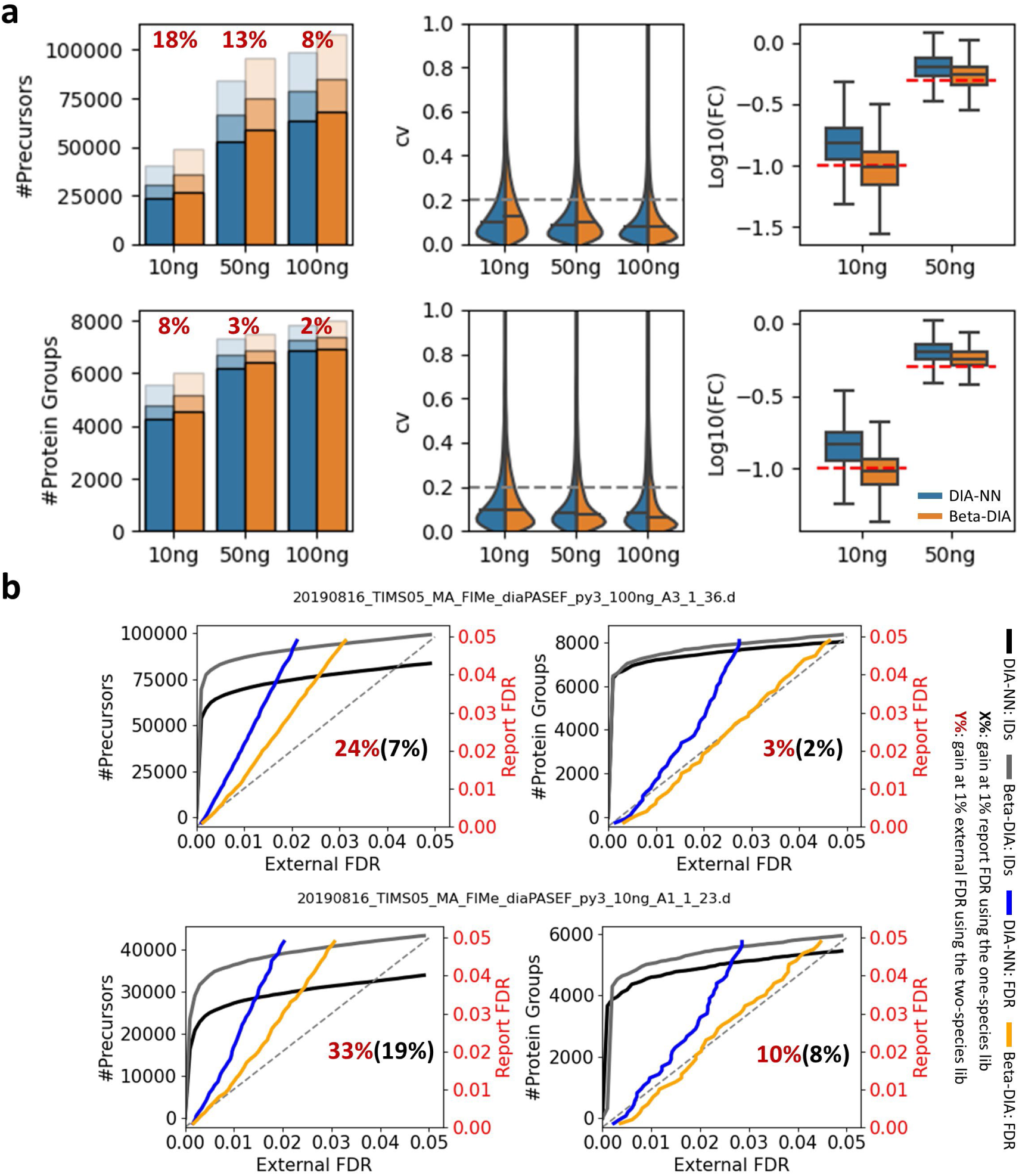
Comparison of the analysis results between Beta-DIA and DIA-NN on the Base-1 dataset. **a)** Qualitative and quantitative results. The bar chart represents identification number from triplicates, with colors ranging from light to dark indicating detection in 1, 2, or all 3 injection replicates. The violin plot represents the coefficients of variation (CV) distributions across three replicates. Then, both the “at least two” method and the average intensity method are used to merge the triplicates. The red numbers above the bars represent the increase achieved by Beta-DIA compared to DIA-NN, calculated by the merged result. The box plot (box denotes the first and third quartiles, the center line shows the median, and the whiskers extend to the most extreme data points within 1.5 times the interquartile range from the box) represents the distribution of fold changes (FC) between lower loading amounts (10 ng, 50 ng) and the maximum loading amount (200 ng). The red dashed line in the box plot represents the expected quantitative ratio. **b)** The FDR validation results using a two-species human-*A. thaliana* spectral library. The two .d files represent the minimum and maximum gains obtained by Beta-DIA using the one-species human spectral library at 1% FDR on precursor level, respectively.

**Supplementary Fig. 8.**
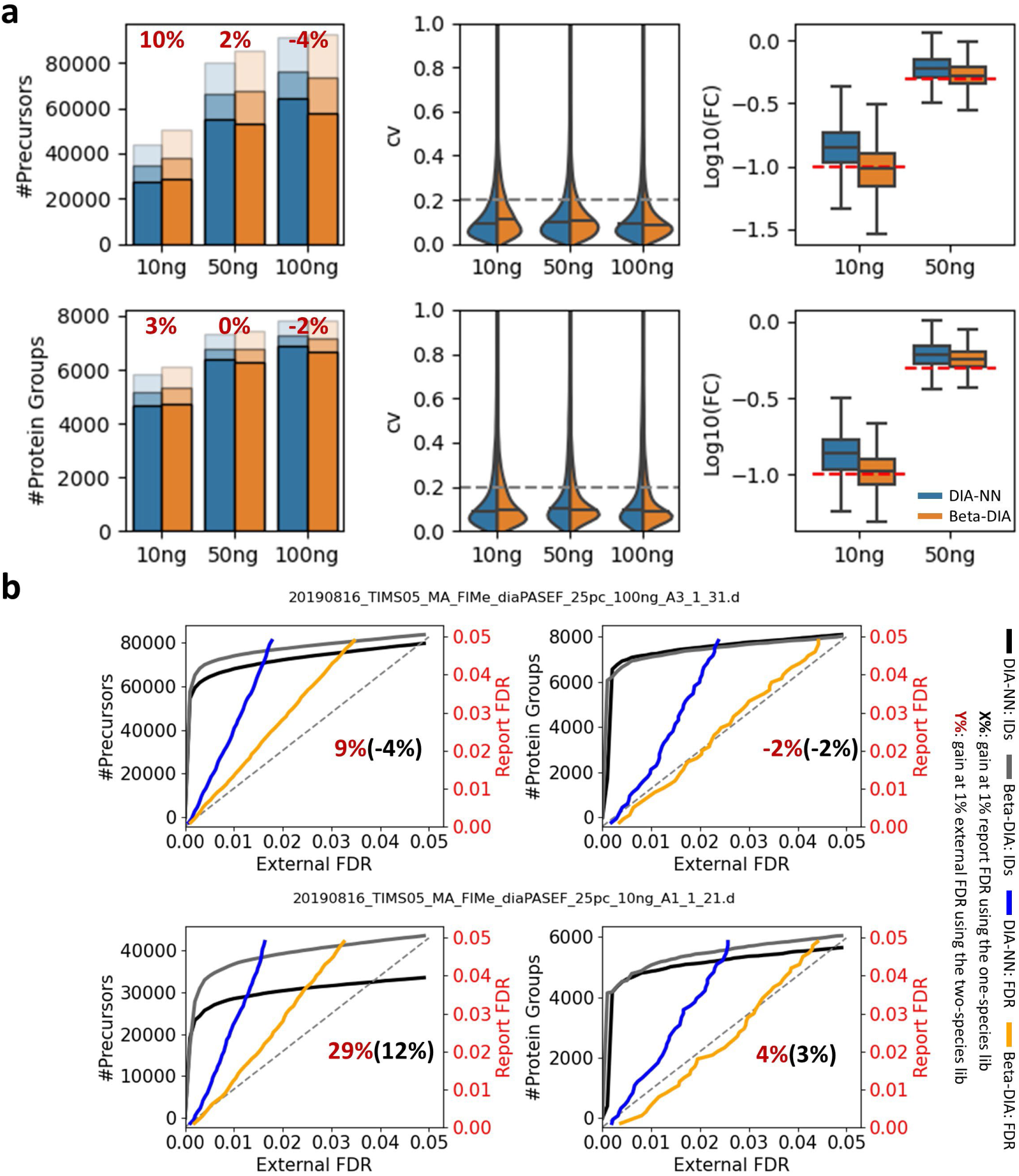
Comparison of the analysis results between Beta-DIA and DIA-NN on the Base-2 dataset. **a)** Qualitative and quantitative results. The bar chart represents identification number from triplicates, with colors ranging from light to dark indicating detection in 1, 2, or all 3 injection replicates. The violin plot represents the coefficients of variation (CV) distributions across three replicates. Then, both the “at least two” method and the average intensity method are used to merge the triplicates. The red numbers above the bars represent the increase achieved by Beta-DIA compared to DIA-NN, calculated by the merged result. The box plot (box denotes the first and third quartiles, the center line shows the median, and the whiskers extend to the most extreme data points within 1.5 times the interquartile range from the box) represents the distribution of fold changes (FC) between lower loading amounts (10 ng, 50 ng) and the maximum loading amount (200 ng). The red dashed line in the box plot represents the expected quantitative ratio. **b)** The FDR validation results using a two-species human-*A. thaliana* spectral library. The two .d files represent the minimum and maximum gains obtained by Beta-DIA using the one-species human spectral library at 1% FDR on precursor level, respectively.

**Supplementary Fig. 9.**
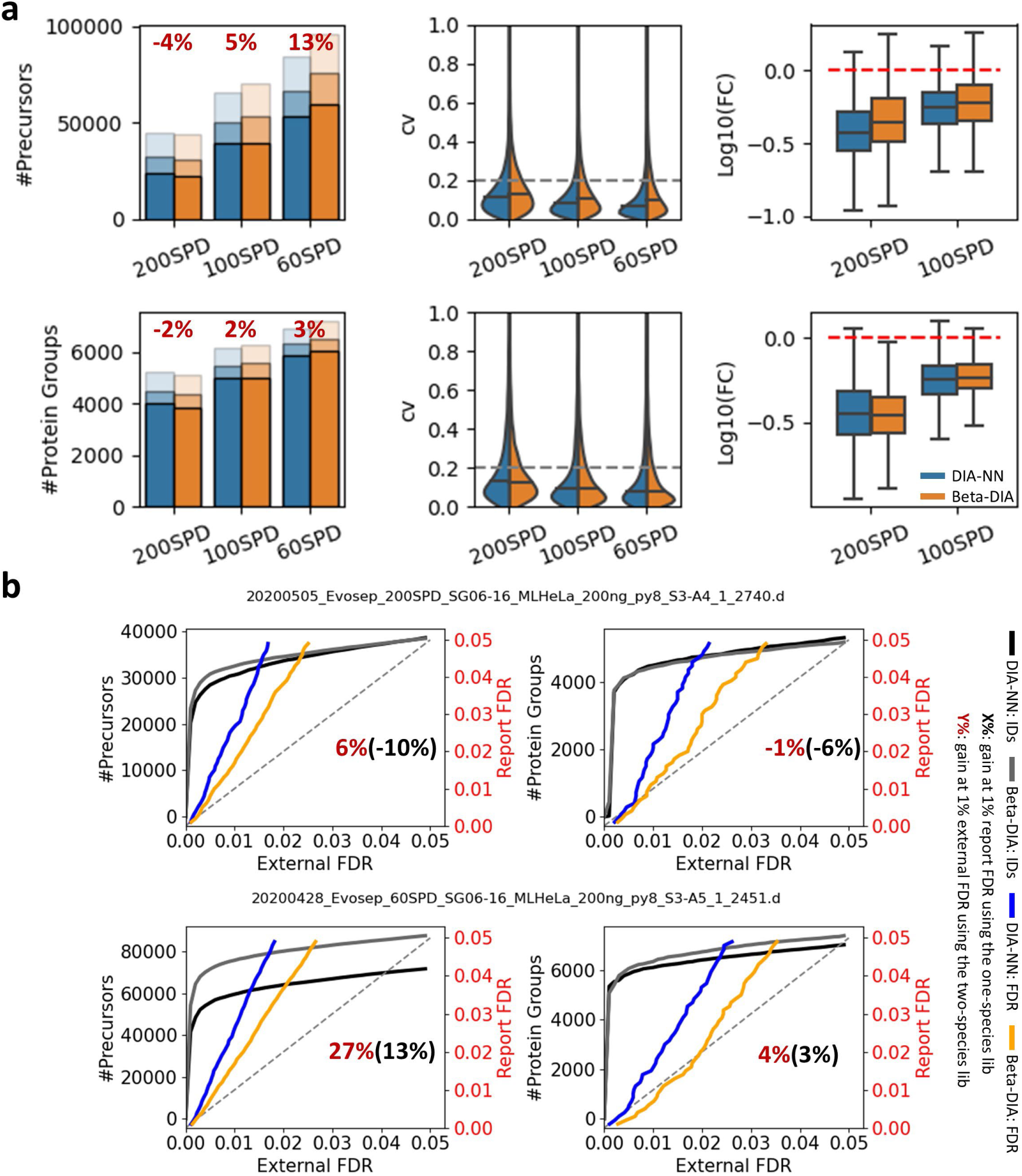
Comparison of the analysis results between Beta-DIA and DIA-NN on the Base-3 dataset. **a)** Qualitative and quantitative results. The bar chart represents identification number from triplicates, with colors ranging from light to dark indicating detection in 1, 2, or all 3 injection replicates. The violin plot represents the coefficients of variation (CV) distributions across three replicates. Then, both the “at least two” method and the average intensity method are used to merge the triplicates. The red numbers above the bars represent the increase achieved by Beta-DIA compared to DIA-NN, calculated by the merged result. The box plot (box denotes the first and third quartiles, the center line shows the median, and the whiskers extend to the most extreme data points within 1.5 times the interquartile range from the box) represents the distribution of fold changes (FC) between the short gradient (200 SPD and 100 SPD, 200 ng) data and the relative long gradient (60 SPD, 200 ng) data. The red dashed line in the box plot represents the expected quantitative ratio. **b)** The FDR validation results using a two-species human-*A. thaliana* spectral library. The two .d files represent the minimum and maximum gains obtained by Beta-DIA using the one-species human spectral library at 1% FDR on precursor level, respectively.

**Supplementary Fig. 10.**
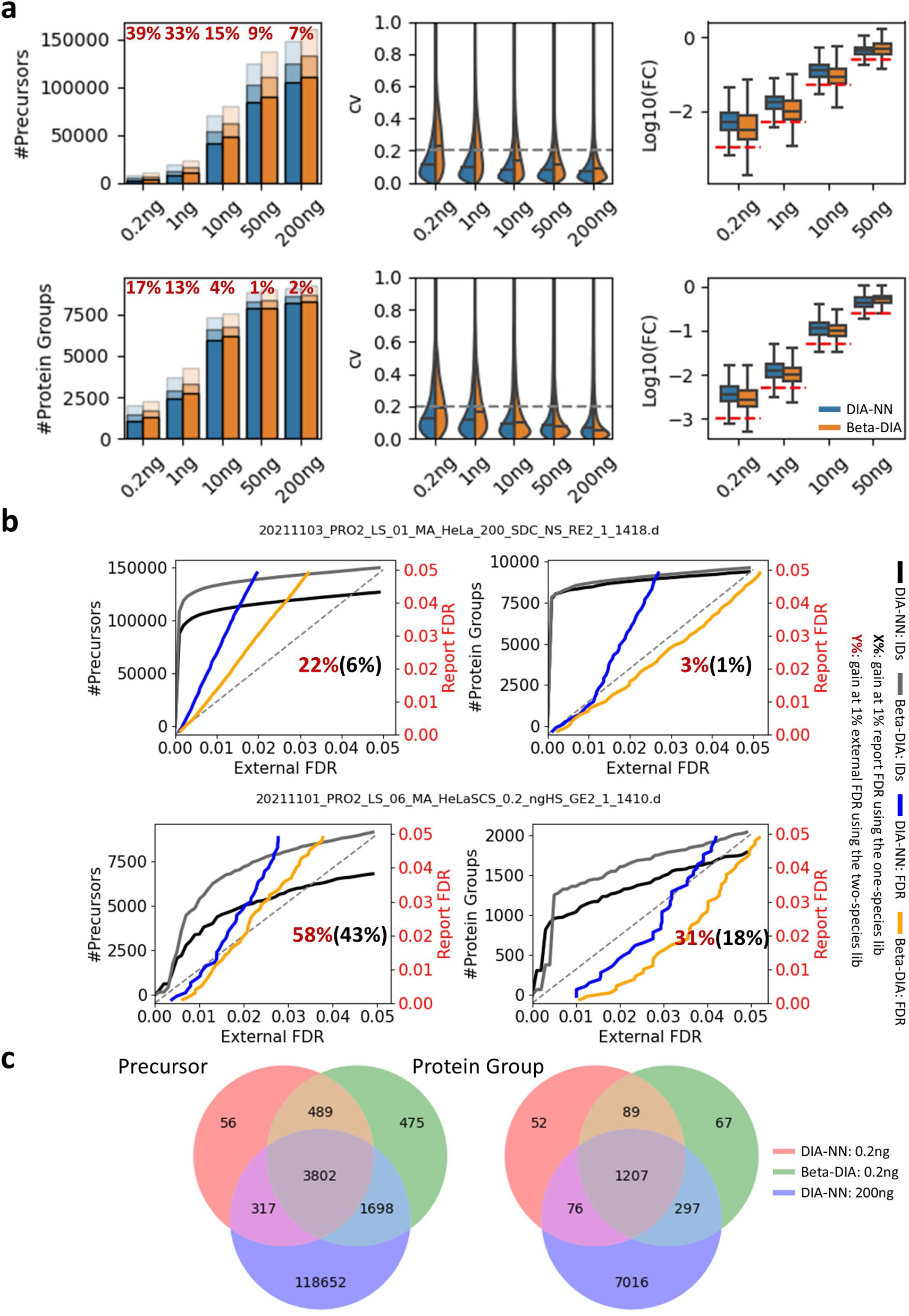
Comparison of the analysis results between Beta-DIA and DIA-NN on the Base-4 dataset. **a)** Qualitative and quantitative results. The bar chart represents identification number from triplicates, with colors ranging from light to dark indicating detection in 1, 2, or all 3 injection replicates. The violin plot represents the coefficients of variation (CV) distributions across three replicates. Then, both the “at least two” method and the average intensity method are used to merge the triplicates. The red numbers above the bars represent the increase achieved by Beta-DIA compared to DIA-NN, calculated by the merged result. The box plot (box denotes the first and third quartiles, the center line shows the median, and the whiskers extend to the most extreme data points within 1.5 times the interquartile range from the box) represents the distribution of fold changes (FC) between lower loading amounts (0.2, 1, 10 and 50 ng) and the maximum loading amount (200 ng). The red dashed line in the box plot represents the expected quantitative ratio. **b)** The FDR validation results using a two-species human-*A. thaliana* spectral library. The two .d files represent the minimum and maximum gains obtained by Beta-DIA using the one-species human spectral library at 1% FDR on precursor level, respectively. **c)** Venn diagrams of the identification results for the lowest (0.2 ng) and highest (200 ng) loading amounts.

**Supplementary Fig. 11.**
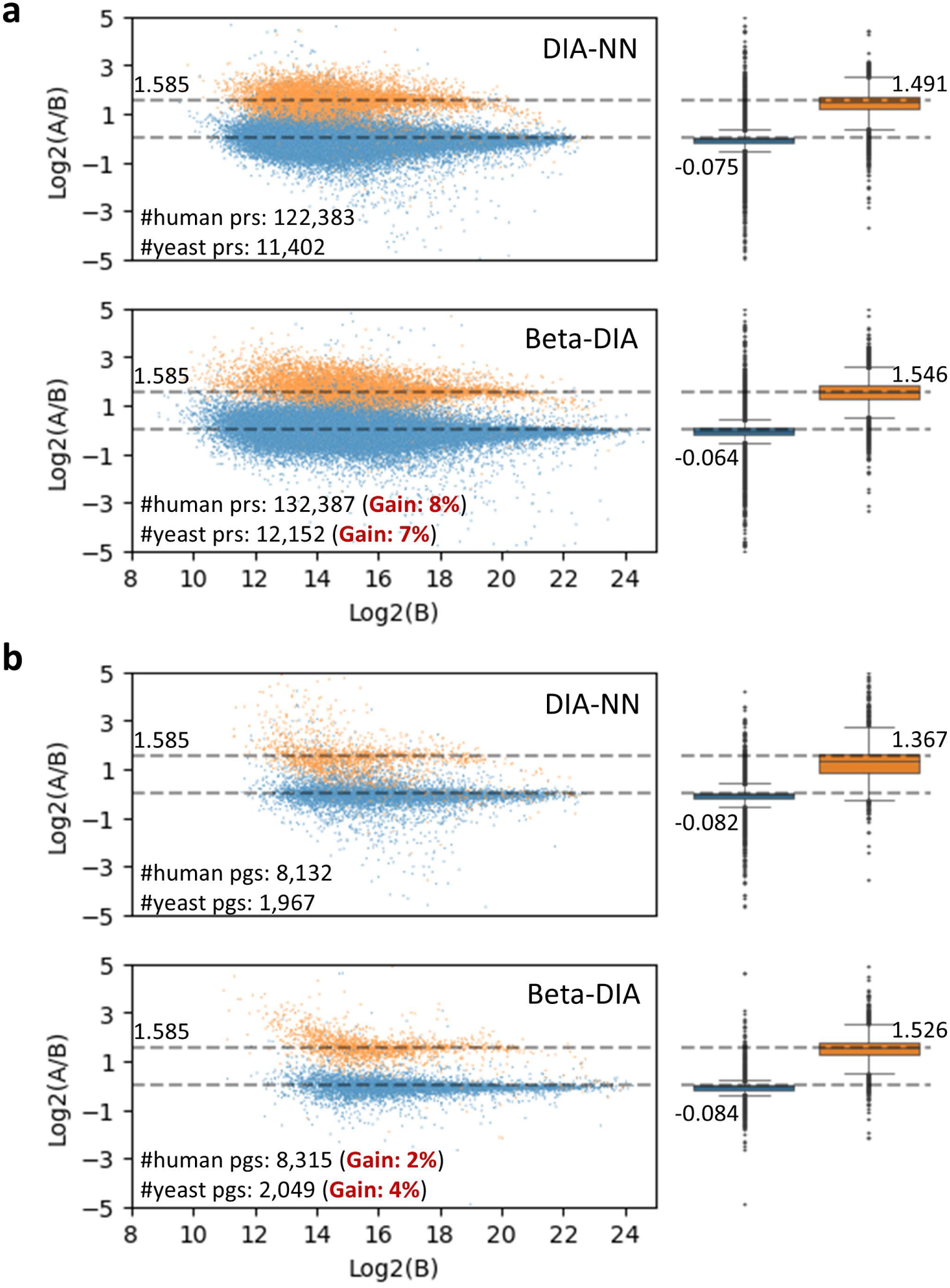
Comparison of the analysis results between Beta-DIA and DIA-NN on the Base-5, a two-proteome mixture dataset. In this dataset, two species (human and yeast) were mixture with different known concentration ratios (human A:B = 1:1, yeast A:B = 45:15). Horizontal dashed lines indicate the expected ratios. On the boxplot, the boxes represent the interquartile range, with the median marked, and the whiskers extend up to 1.5x the interquartile range. **a)** on precursor level. **b)** on protein group level.

**Supplementary Fig. 12.**
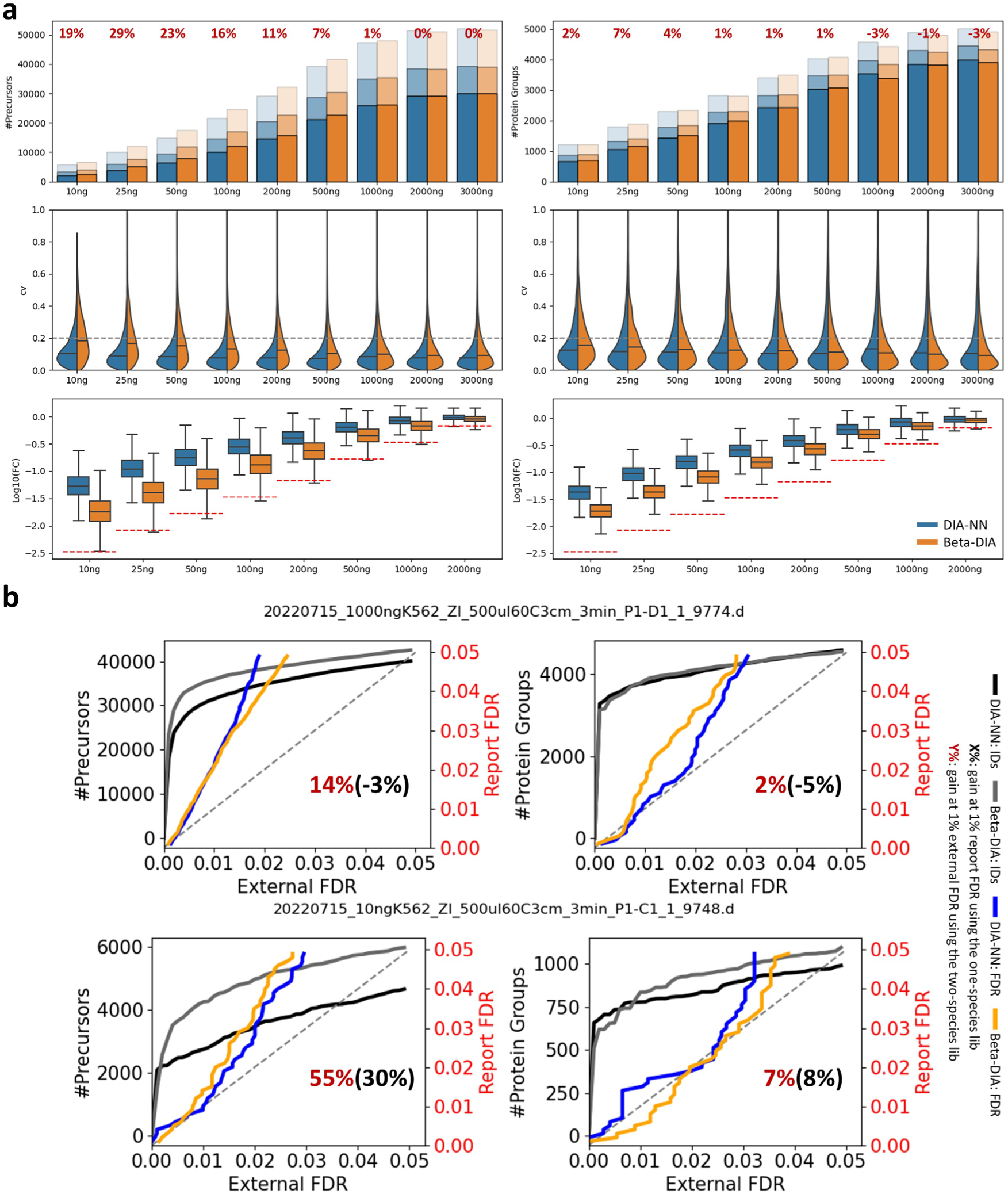
Comparison of the analysis results between Beta-DIA and DIA-NN on the Extra-1-1 dataset. **a)** Qualitative and quantitative results. The bar chart represents identification number from triplicates, with colors ranging from light to dark indicating detection in 1, 2, or all 3 injection replicates. The violin plot represents the coefficients of variation (CV) distributions across three replicates. Then, both the “at least two” method and the average intensity method are used to merge the triplicates. The red numbers above the bars represent the increase achieved by Beta-DIA compared to DIA-NN, calculated by the merged result. The box plot (box denotes the first and third quartiles, the center line shows the median, and the whiskers extend to the most extreme data points within 1.5 times the interquartile range from the box) represents the distribution of fold changes (FC) between lower loading amounts (10-2000ng) and the maximum loading amount (3000 ng). The red dashed line in the box plot represents the expected quantitative ratio. **b)** The FDR validation results using a two-species human-*A. thaliana* spectral library. The two .d files represent the minimum and maximum gains obtained by Beta-DIA using the one-species human spectral library at 1% FDR on precursor level, respectively.

**Supplementary Fig. 13.**
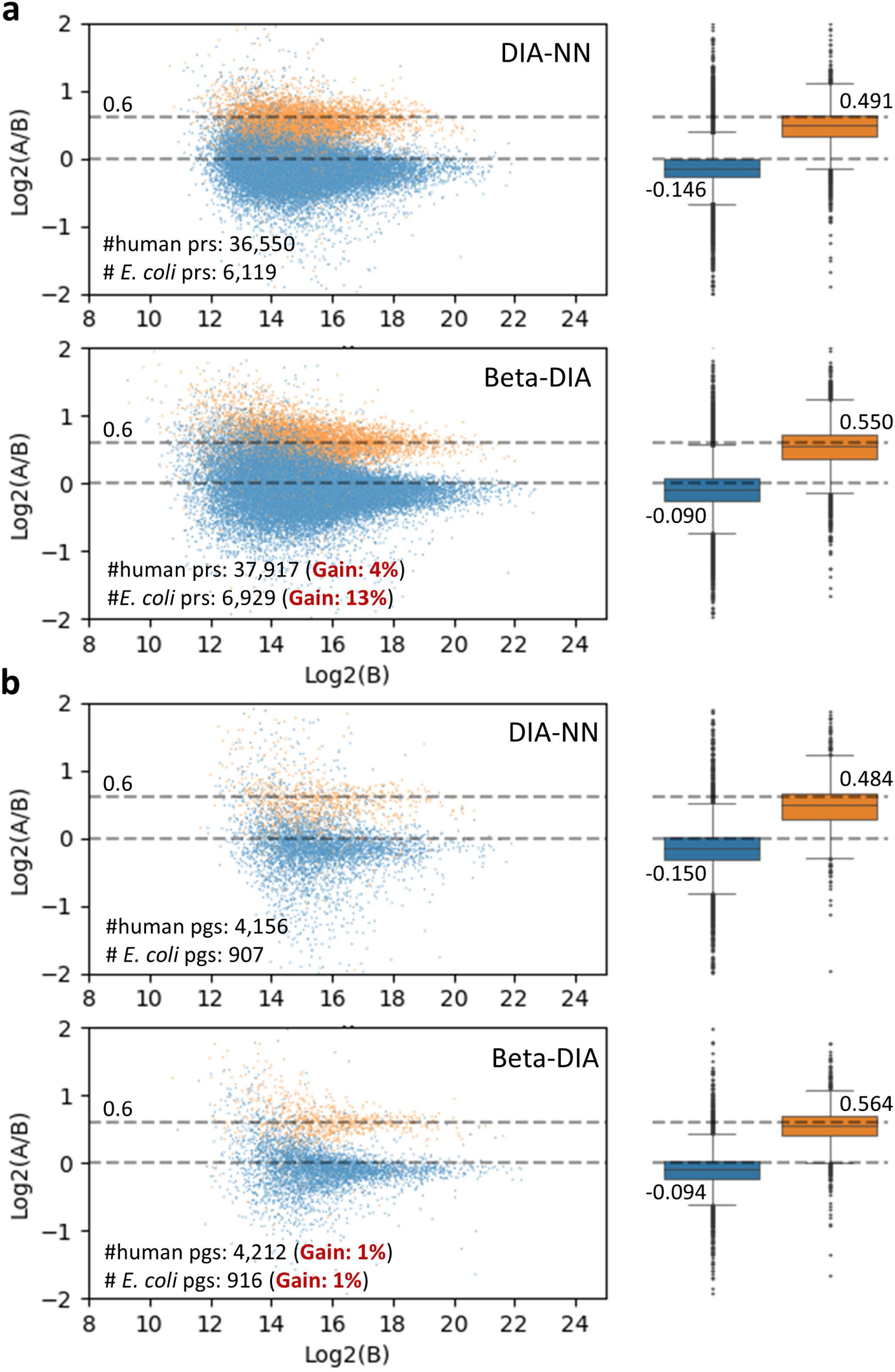
Comparison of the analysis results between Beta-DIA and DIA-NN on the Extra-1-2, a two-proteome mixture dataset. In this dataset, two species (human and *E. coli*) were mixture with different known concentration ratios (human A:B = 1:1, *E. coli* A:B = 50:33). Horizontal dashed lines indicate the expected ratios. On the boxplot, the boxes represent the interquartile range, with the median marked, and the whiskers extend up to 1.5x the interquartile range. **a)** on precursor level. **b)** on protein group level.

**Supplementary Fig. 14.**
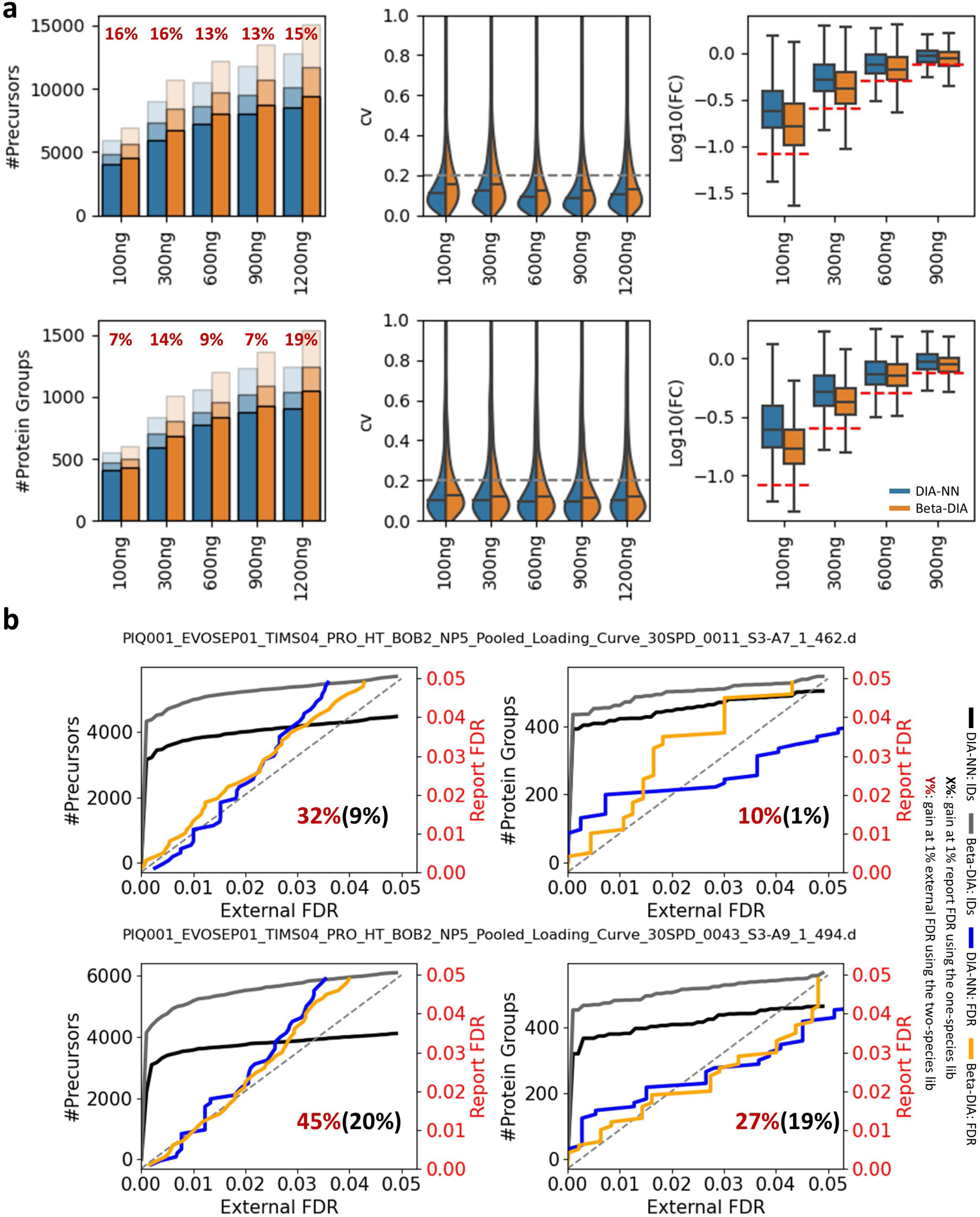
Comparison of the analysis results between Beta-DIA and DIA-NN on the Extra-2 dataset. **a)** Qualitative and quantitative results. The bar chart represents identification number from triplicates, with colors ranging from light to dark indicating detection in 1, 2, or all 3 injection replicates. The violin plot represents the coefficients of variation (CV) distributions across three replicates. Then, both the “at least two” method and the average intensity method are used to merge the triplicates. The red numbers above the bars represent the increase achieved by Beta-DIA compared to DIA-NN, calculated by the merged result. The box plot (box denotes the first and third quartiles, the center line shows the median, and the whiskers extend to the most extreme data points within 1.5 times the interquartile range from the box) represents the distribution of fold changes (FC) between lower loading amounts (100-900 ng) and the maximum loading amount (1200 ng). The red dashed line in the box plot represents the expected quantitative ratio. **b)** The FDR validation results using a two-species human-*A. thaliana* spectral library. The two .d files represent the minimum and maximum gains obtained by Beta-DIA using the one-species human spectral library at 1% FDR on precursor level, respectively.

**Supplementary Fig. 15.**
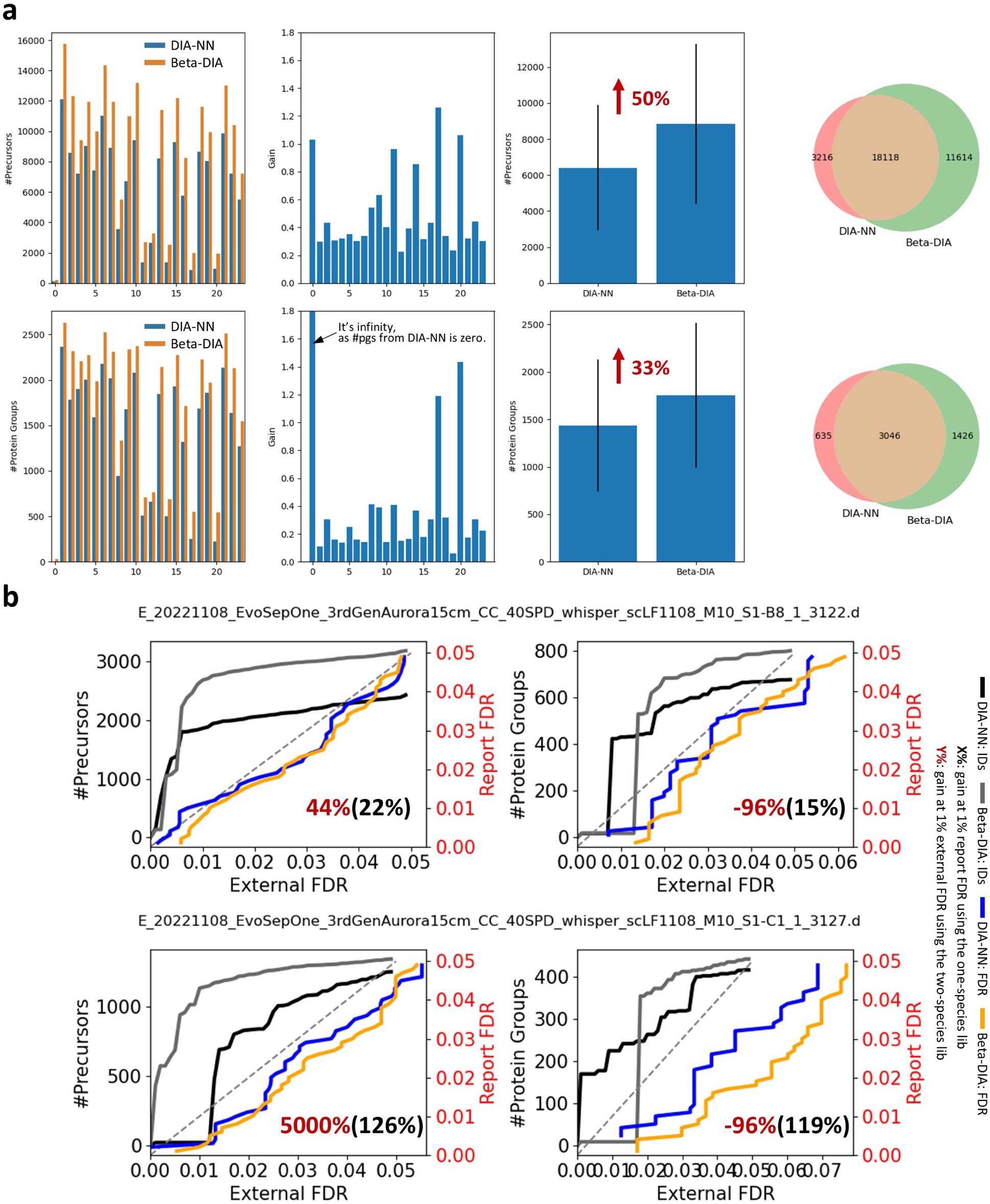
Comparison of the analysis results between Beta-DIA and DIA-NN on the Extra-3-1 dataset. **a)** Qualitative results. Starting from the left, the first bar chart shows the number of identifications for different single-cell samples; the second bar chart displays the gains achieved by Beta-DIA compared to DIA-NN; and the third bar chart presents the average number of identifications along with the standard deviation. The Venn diagram on the far right represents the identification results from all single-cell samples. **b)** The FDR validation results using a two-species human-*A. thaliana* spectral library. The two .d files represent the minimum and maximum gains obtained by Beta-DIA using the one-species human spectral library at 1% FDR on precursor level, respectively.

**Supplementary Fig. 16.**
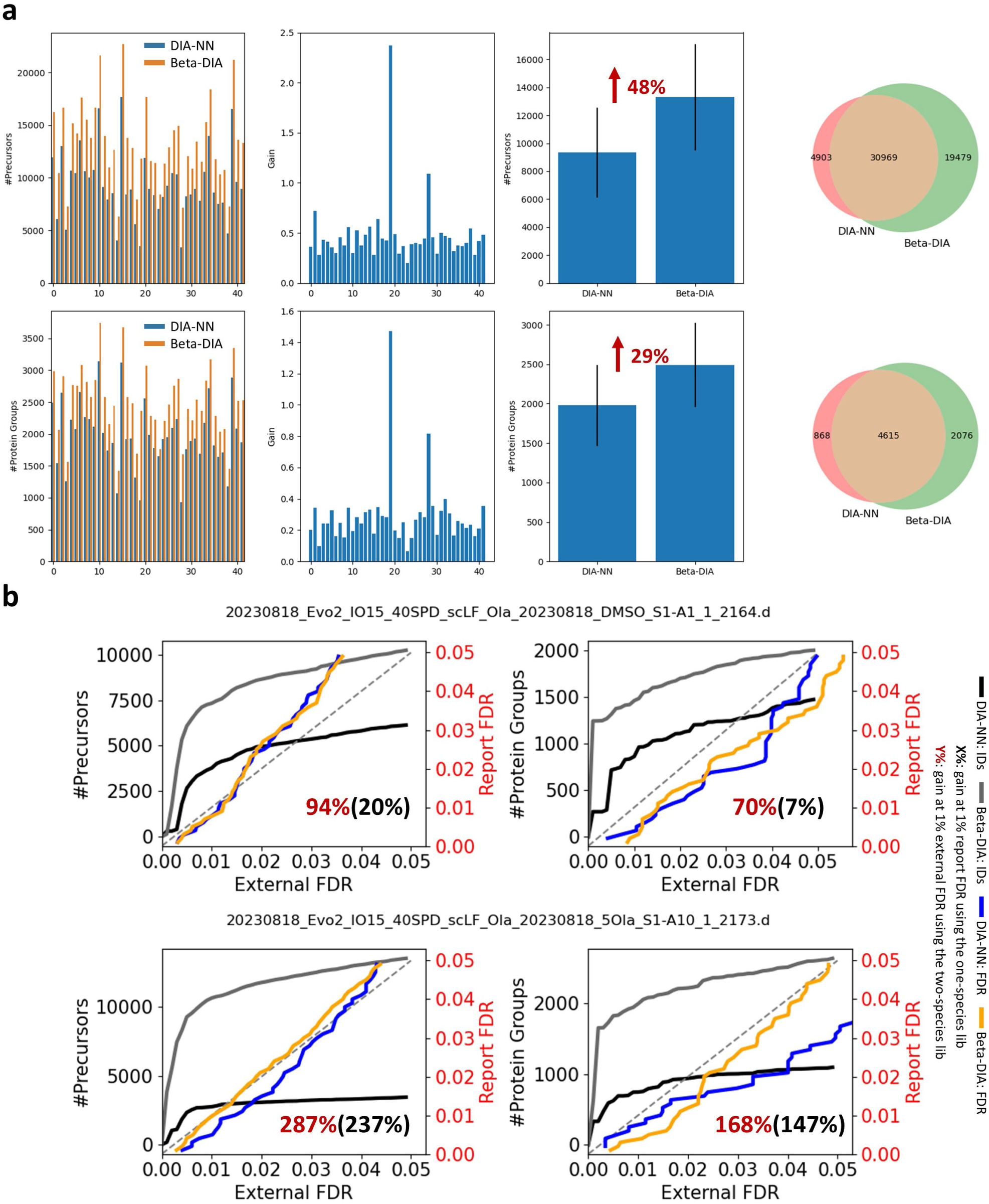
Comparison of the analysis results between Beta-DIA and DIA-NN on the Extra-3-2 dataset. **a)** Qualitative results. Starting from the left, the first bar chart shows the number of identifications for different single-cell samples; the second bar chart displays the gains achieved by Beta-DIA compared to DIA-NN; and the third bar chart presents the average number of identifications along with the standard deviation. The Venn diagram on the far right represents the identification results from all single-cell samples. **b)** The FDR validation results using a two-species human-*A. thaliana* spectral library. The two .d files represent the minimum and maximum gains obtained by Beta-DIA using the one-species human spectral library at 1% FDR on precursor level, respectively.

